# Histone chaperone Nucleophosmin regulates transcription of key genes involved in oral tumorigenesis

**DOI:** 10.1101/852095

**Authors:** Parijat Senapati, Suchismita Dey, Deepthi Sudarshan, Aditya Bhattacharya, Shyla G, Sadhan Das, Surabhi Sudevan, Tessy Thomas Maliekal, Tapas K. Kundu

## Abstract

Nucleophosmin (NPM1) is a multifunctional histone chaperone that can activate RNA Polymerase II-driven chromatin transcription. Acetylation of NPM1 by acetyltransferase p300 has been shown to further enhance its transcription activation potential. Moreover, its total and acetylated pools are increased in oral squamous cell carcinoma. However, the role of NPM1 or its acetylated form (AcNPM1) in transcriptional regulation in cells is not fully elucidated. Using ChIP-seq analyses, we show that AcNPM1 co-occupies marks of active transcription at promoters and DNase I hypersensitive sites at enhancers. Moreover, NPM1 interacts with proteins involved in transcription, including RNA Pol II, general transcription factors, mediator subunits, histone acetyltransferase complexes, and chromatin remodelers. Moreover, its histone chaperone also contributes to transcriptional activation. We further show that AcNPM1 regulates key genes required for proliferation, migration and invasion potential of oral cancer cells and knockdown of NPM1 mitigates these processes in cells as well as orthotopic tumors in mice. Collectively, these results establish that AcNPM1 functions as a coactivator and regulates the expression of key genes involved in oral tumorigenesis.

## INTRODUCTION

Nucleophosmin (NPM1/B23), is a ubiquitously expressed histone chaperone (Okuwaki et al. 2001; Swaminathan et al. 2005) that belongs to the Nucleoplasmin family of histone chaperones (Lee et al. 2007). As the name suggests, histone chaperone proteins bind to histones and assist in nucleosome assembly and disassembly processes during DNA replication, transcription and repair (Gurard-Levin et al. 2014). NPM1 is mainly localized in the nucleolus, however, it is a highly dynamic nucleo-cytoplasmic shuttling protein with established functions in the nucleolus, nucleoplasm, and cytoplasm (Lindström 2011). Some of its well-characterized functions include regulation of centrosome duplication (Okuda 2002), ribosome biogenesis (Savkur and Olson 1998), nuclear import and molecular chaperone activity (Szebeni and Olson 1999), as well as induction of p300 autoacetylation (Arif et al. 2010; Kaypee et al. 2018). In addition, NPM1 function is regulated by post-translational modifications including acetylation, phosphorylation, SUMOylation, and ubiquitination (Colombo et al. 2011). Being a histone chaperone, NPM1 has roles in DNA replication, repair as well as transcription. NPM1 regulates RNA Polymerase I-mediated transcription through its histone chaperone activity (Murano et al. 2008). Moreover, it binds with histones and enhances acetylation-dependent *in vitro* transcription from a chromatinized template. Acetylation by p300 enhances its binding affinity for histones as well as its histone chaperone activity (Swaminathan et al. 2005). Acetylation also augments its ability to activate acetylation-dependent chromatin transcription over and above unmodified NPM1 presumably due to the enhanced nucleosome disassembly activity of acetylated NPM1 (AcNPM1) (Swaminathan et al. 2005). Furthermore, the AcNPM1 pool localizes to the nucleoplasm and colocalizes with RNA Polymerase II (Pol II) (Shandilya et al. 2009). NPM1 has also been shown to regulate the expression of specific genes by binding to transcription factors including p53 (Colombo et al. 2002), NF-κB (Dhar et al. 2004), YY1 (Inouye and Seto 1994), c-myc (Li et al. 2008) and IRF1 (Kondo et al. 1997). However, the transcriptional role of AcNPM1 at the genome-wide level is not known.

In addition, NPM1 has been proposed to have an oncogenic function as it is overexpressed in tumors of diverse histological origins as well as in hematological malignancies (Grisendi et al. 2006). NPM1 is a direct transcriptional target of oncogenic transcription factors c-myc (Zeller et al. 2001), c-fos as well as a gain-of-function mutant of p53 (Senapati et al. 2018). We have earlier shown that NPM1 is overexpressed in oral squamous cell carcinoma (OSCC) and AcNPM1 levels increase with advancing tumor grade (Shandilya et al. 2009). NPM1 exists as an oligomer in cells, a property imparted by its highly conserved N-terminal domain that is a characteristic feature shared by the Nucleoplasmin family of histone chaperones (Frehlick et al. 2007). Oligomerization has been earlier shown to be important for its molecular chaperone activity (Hingorani et al. 2000) as well as the induction of p300 autoacetylation (Kaypee et al. 2018). Interestingly, small molecules and RNA aptamers that target NPM1 oligomerization have been reported to be effective in inducing apoptosis in cancer cells (Qi et al. 2008; Jian et al. 2009). These studies indicate that NPM1 overexpression is causally connected with the maintenance of the proliferative state of tumor cells. Given the role of NPM1 as a histone chaperone, we postulated that NPM1/AcNPM1 is involved in transcriptional regulation of genes and regulates key gene networks essential for oral tumor growth.

We report here an investigation into the role of AcNPM1 in regulating transcription and the functional consequence of knocking down NPM1 on OSCC tumor growth. We report the first genome-wide profile of AcNPM1 occupancy and show that AcNPM1 is associated with active transcription. AcNPM1 occupies transcriptional regulatory elements such as promoters and enhancers through its interaction with RNA Pol II, general transcription factors (GTFs), transcription factors as well as histone acetyltransferases and chromatin remodeling complexes. We further show that its histone chaperone activity is crucial for its transcriptional activation potential. In oral cancer cells, AcNPM1 regulates key pathways associated with cell proliferation, migration, invasion, and NPM1 knockdown attenuates oral tumor growth in mouse xenografts. In summary, our results provide a comprehensive understanding of the RNA Pol II-driven transcriptional regulation function of NPM1 and its consequences in oral tumorigenesis.

## RESULTS

### AcNPM1 is enriched at active gene promoters

We have previously established that the acetylated NPM1 pool (AcNPM1) localizes to the nucleoplasm in contrast to the nucleolar localization of NPM1 (Shandilya et al. 2009). To directly interrogate the gene targets and genome-wide occupancy of AcNPM1, we performed chromatin immunoprecipitation sequencing (ChIP-seq) using an in-house generated polyclonal antibody against NPM1 acetylated at residues K229 and K230 (see Methods). We tested the specificity of the antibody using western blots, immunoprecipitation (IP) and dot-blot assays. The antibody specifically detected the acetylated form and not the unacetylated form of NPM1 (Figure S1A). Further, the antibody was able to pull down NPM1 from HeLa S3 lysates (Figure S1B). The antibody also showed the characteristic nucleoplasmic localization of AcNPM1 as has been reported earlier (Figure S1C) (Shandilya et al. 2009). In addition, dot-blot assays further confirmed that the antibody specifically detects the acetylation at residues K229 and K230 as it did not detect NPM1 peptides acetylated at other residues, the unacetylated peptide or the unmodified recombinant NPM1 protein (Figure S1D). The AcNPM1 antibody also did not detect acetylation at specific histone residues (Figure S1E). Similarly, the AcNPM1 antibody could be effectively blocked only with NPM1(K229, K230)ac peptide and not any other acetylated or unacetylated peptides (Figure S1F and G). These results show that the AcNPM1 antibody is highly specific to acetylation of K229 and K230 residues of NPM1.

We performed ChIP-seq in HeLa S3 cells to interrogate the genome-wide occupancy of AcNPM1 (see Methods). We obtained more than 82 million single-end reads from each replicate of ChIP and input (Table S1). Sequencing adapters were removed, and reads were aligned to hg19 assembly of the human genome using bowtie2 (see Methods). We called broad peaks using MACS2 (Zhang et al. 2008) and identified 24660 peaks conserved between the 2 replicates. We compared the genomic location of the AcNPM1 peaks in relation to the transcription start sites (TSS) of hg19 RefSeq genes. About 40.65% (10056) of the AcNPM1 peaks were located within 1kb of TSS and about 60.1% (14867) of all peaks were within 10kb of the TSS indicating enrichment of AcNPM1 at TSS or at promoter-proximal regions (Figure 1A). In comparison, a random selection of peaks of the same size on the human genome showed only 16.48% of the peaks at promoter-proximal regions (Figure 1A). Genome browser view of a section of the chromosome 19 shows AcNPM1 peaks at TSS of multiple genes (Figure 1B). Heatmap of ChIP-seq tag density at TSS ± 2kb of all RefSeq genes also shows the enrichment of AcNPM1 at promoter regions (Figure 1C). We next compared the AcNPM1 peaks to genome-wide profiles of other histone modifications in HeLa S3 cells available from the ENCODE consortium (Consortium 2012). Jaccard index, a measure of similarity or overlap, was calculated for pairwise comparisons of AcNPM1 with active (H3K4me3, H3K9ac, H3K27ac, H3K4me1, H3K4me2, H3K79me2, H3K36me3, and H4K20me1) and repressive (H3K9me3 and H3K27me3) histone modification marks. The Jaccard indexes were plotted as a heatmap with hierarchical clustering based on similarity. AcNPM1 clustered strongly with known active promoter marks such as H3K27ac, H3K9ac, H3K4me3, and moderately with H3K4me1 and H3K4me2 that are also observed at promoters (Figure 1D). In contrast, AcNPM1 showed no overlap with other active histone modifications that are enriched on gene bodies namely H3K79me2, H3K36me3 and H4K20me1 as well as repressive histone modifications H3K9me3 and H3K27me3 (Figure 1D). These results show that AcNPM1 occupies gene promoters with active histone modifications.

**Figure 1:**
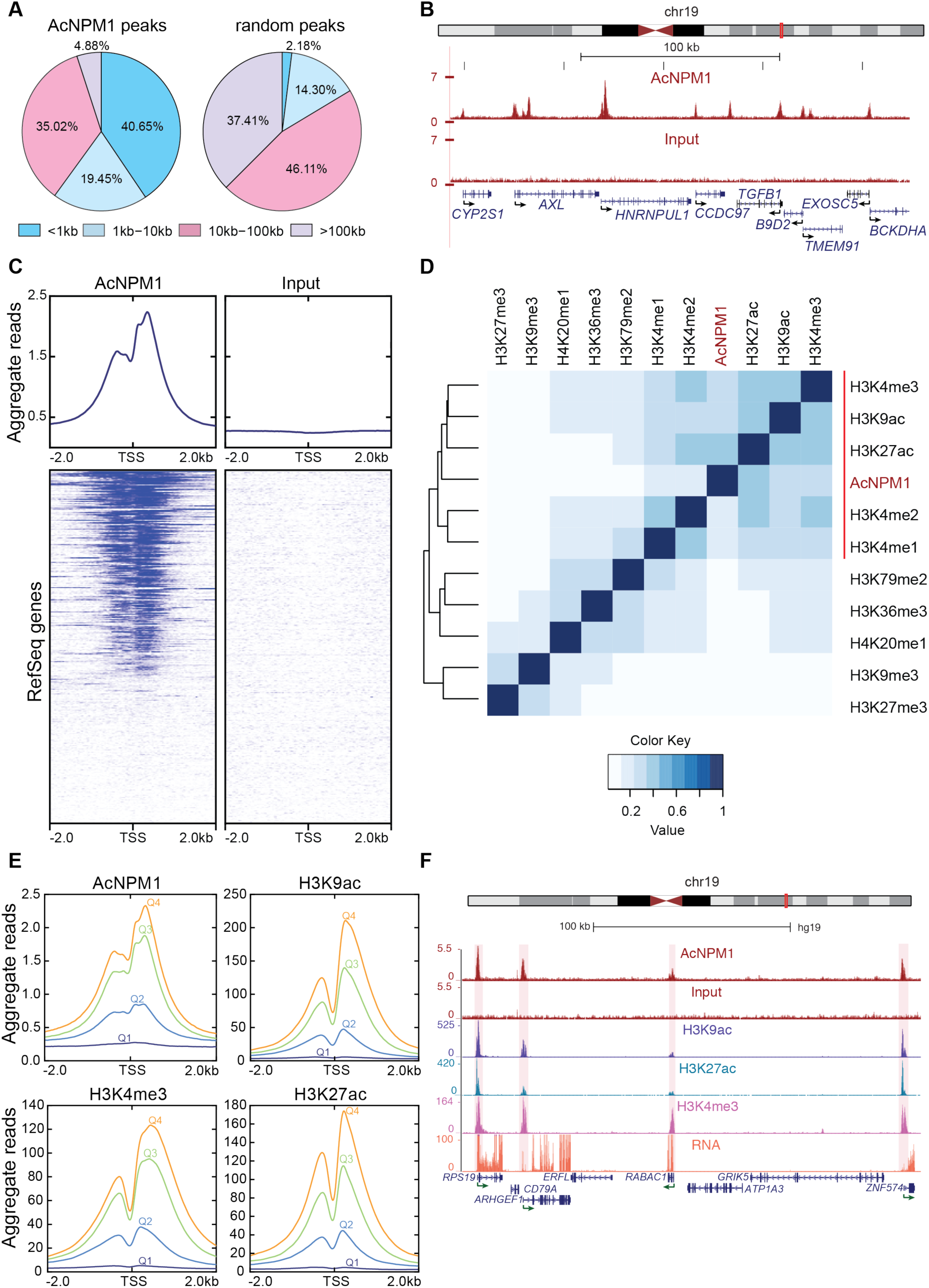
AcNPM1 is enriched at active gene promoters. (A) Genomic distribution of AcNPM1 ChIP-seq peaks plotted according to their distance from RefSeq TSS (left pie chart). A random distribution of peaks of similar size (expected) is shown for comparison (right). (B) UCSC genome browser snapshot showing the distribution of normalized reads for AcNPM1 and Input at the TSS of indicated genes on chr19 (http://genome.ucsc.edu/). (C) Heatmaps of normalized reads for AcNPM1 and Input at TSS ± 2kb regions of all RefSeq genes. Aggregate profile of AcNPM1 and Input read density is shown above the heatmaps. (D) Clustered heatmap showing the Jaccard indexes for overlap between peaks for AcNPM1 and indicated histone modification marks. The color scale for Jaccard indexes is shown. (E) Normalized read counts for AcNPM1 and other histone modification marks as indicated, are plotted in 10bp bins in a TSS ± 2kb window for all RefSeq transcripts. Q1 (dark blue) and Q4 (orange) represent the quartile of genes with the lowest and highest level of expression respectively. (F) UCSC genome browser snapshots showing the distribution of normalized reads for AcNPM1, Input, histone modification marks H3K27ac, H3K4me3, and H3K9ac at the TSS of indicated genes as well as an RNA track showing their expression levels.

To test whether AcNPM1 is associated with active transcription, RefSeq transcripts were divided into four quartiles based on their expression levels in HeLa S3 cells. Highly expressed transcripts showed higher enrichment of AcNPM1 at their promoters as compared to those that showed lower expression levels (Figure 1E). Moreover, AcNPM1 was absent at genes with no detectable expression in HeLa S3 cells (Figure 1E). Other promoter enriched marks, H3K27ac, H3K9ac, and H3K4me3 were also plotted and showed the expected enrichment on these highly transcribed genes (Figure 1E). Heatmaps of AcNPM1 ChIP-seq tags on promoters (TSS ± 2kb) of genes sorted by expression levels in HeLa S3 cells also confirm that AcNPM1 is enriched at promoters of transcriptionally active genes (Figure S2) along with H3K27ac, H3K9ac, and H3K4me3. Genome browser screenshots further show the co-occupancy of AcNPM1 with H3K27ac, H3K9ac, and H3K4me3 peaks at promoters of expressed genes (Figure 1F). These results indicate that AcNPM1 is associated with active transcription at gene promoters.

### AcNPM1 co-occupies with RNA Pol II, chromatin remodeling factors and TFs at transcriptional regulatory elements

In order to further characterize the genomic localization of AcNPM1, we compared AcNPM1 peaks in HeLa S3 cells with the combined segmentation (ChromHMM + Segway) (Hoffman et al. 2013) of the HeLa S3 genome from UCSC genome browser (Kent et al. 2002). 41.6% of AcNPM1 peaks overlapped with TSS regions and 25.7% of them overlapped with known enhancer regions in the HeLa S3 genome (Figure 2A). Interestingly, AcNPM1 peaks also overlapped with CTCF-bound regions (Figure 2A), in agreement with an earlier report that showed NPM1 and CTCF co-binding at insulator elements (Yusufzai et al. 2004). We also compared the extent of occupancy of known TSS and enhancer classifications by AcNPM1 in HeLa S3 cells. We found that ∼90% of the TSS as classified by the combined segmentation classifications (ChromHMM + Segway) were bound by AcNPM1 and ∼54% of the enhancers were bound by AcNPM1 (Figure 2B). Genome browser views confirm the presence of AcNPM1 peaks on TSS and enhancer regions (Figure 2C). These results indicate that AcNPM1 is associated with active transcriptional regulatory elements. We further examined the AcNPM1 signal at DNase I hypersensitive sites (DHS), which is another measure of active regulatory elements. AcNPM1 ChIP signal is significantly higher at DHS regions as compared to non-DHS regions (Figure 2D). Similarly, we compared AcNPM1 ChIP signal at H3K27ac, p300, and RNA Pol II peaks that are other markers of active regulatory elements. AcNPM1 was highly enriched in regions with high H3K27ac levels (Figure 2E), and regions that overlapped with p300 (Figure 2F) and RNA Pol II (Figure 2G). These results show that AcNPM1 is associated with active transcription usually found at regulatory elements such as promoters and enhancers. To further understand the mechanism of AcNPM1 occupancy at active regulatory elements, we identified transcription factor binding motifs enriched at AcNPM1 peaks. We found enrichment of several transcription factor (TF) binding motifs from diverse families such as ETS, E2F, IRF, bZip, bHLH, STAT, RFX, and Zinc fingers (Figure 2H). In addition, to identify the factors co-bound at AcNPM1 peaks, we compared the binding profiles for several TFs and chromatin-related factors in HeLa S3 cells available from the ENCODE project (Consortium 2012), at AcNPM1 peaks. Only proteins that overlapped with at least 10% of AcNPM1 peaks were selected for the comparison. The transcription factor MAX has the highest co-occupancy at AcNPM1 peaks (Figure 2I). About 60% of AcNPM1 peaks are co-bound by MAX followed by RNA Pol II and Chromodomain Helicase DNA Binding Protein 2 (CHD2) (Figure 2I). For an unbiased view of co-binding factors, we plotted the correlation values for co-occupancy of different factors with AcNPM1 as a heatmap with hierarchical clustering to show the co-bound factors (Figure 2J). The largest cluster of co-bound factors contained related TFs MYC, MAX, MAZ, MXI1 along with TAF_II_250, CHD2, and RNA Pol II (Figure 2J). A second cluster co-bound by c-fos, c-jun, JUND, p300 and STAT3 was also observed (Figure 2J). Other significant clusters included AcNPM1 peaks co-bound by ELK1, ELK4, and GABPA; E2F4, E2F6, and E2F1; AP2alpha and AP2gamma; NFYA and NFYB; CTCF, RAD21, and SMC3 (Figure 2J). We earlier observed about 6% of the AcNPM1 peaks are at CTCF bound regions (Figure 2A). CTCF, RAD21, and SMC3 are present in a complex at boundary elements (Parelho et al. 2008; Wendt et al. 2008). Interestingly, binding sites for NRF1, E2F family, JUN, AP-1, CTCF, STAT family, ELK family were also identified to be enriched in AcNPM1 peaks (Figure 2H). Together, these results show that AcNPM1 co-occupies with RNA Pol II, transcription factors and chromatin remodeling factors at transcriptional regulatory elements.

**Figure 2:**
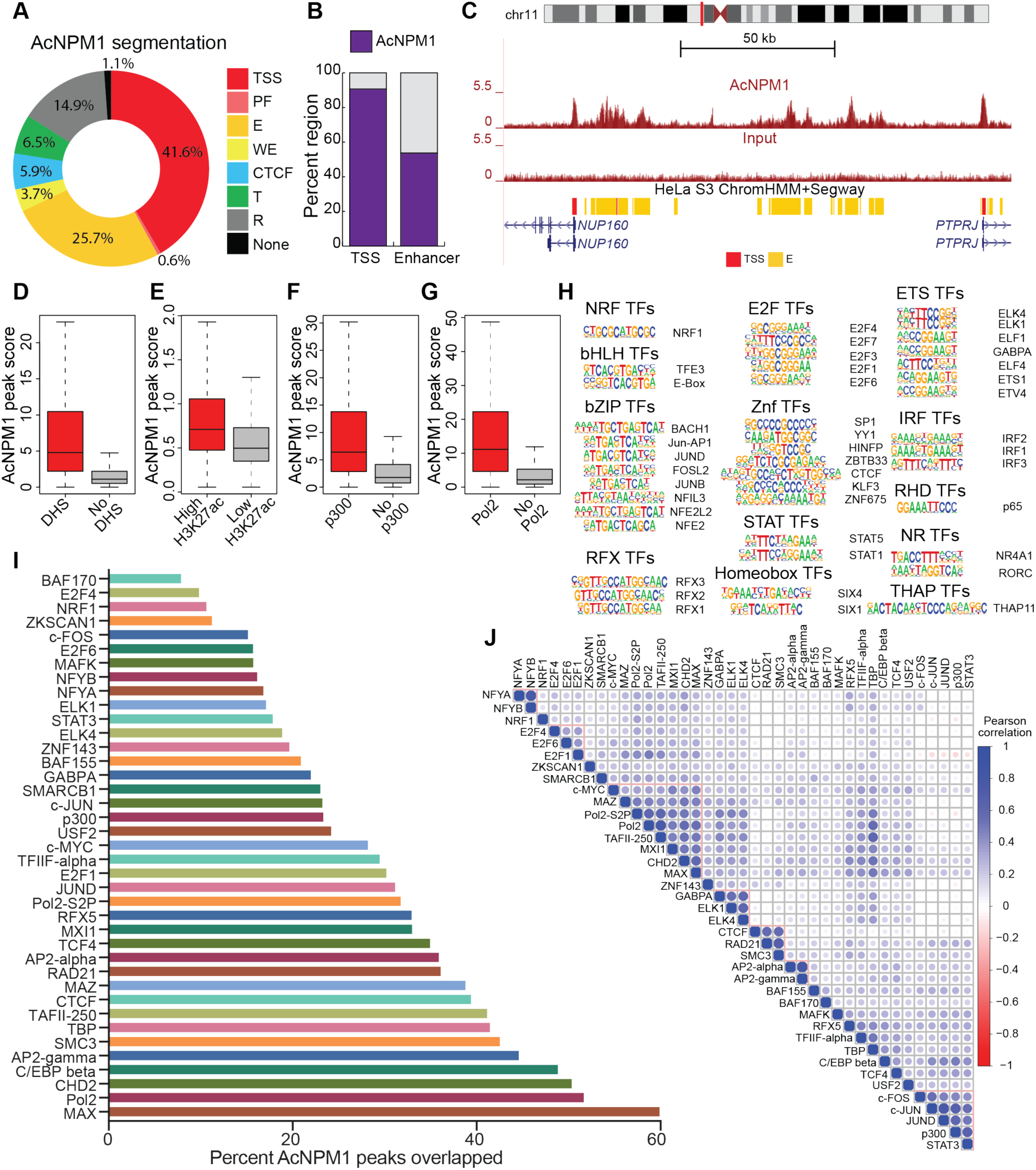
AcNPM1 co-occupies with RNA Pol II, chromatin remodeling factors and transcription factors at transcriptional regulatory elements. (A) Plot showing the percent number of AcNPM1 peaks overlapped with ChromHMM + Segway combined segmentation for HeLa S3 genome from the UCSC genome browser. (Key: TSS, predicted promoter region including TSS; PF, predicted promoter flanking region; E, enhancer; WE, predicted weak enhancer or open chromatin cis-regulatory element; CTCF, CTCF enriched element; T, predicted transcribed region; R, predicted repressed or low activity region; None, unclassified). (B) Percent number of TSS and enhancer regions identified by ChromHMM + Segway combined segmentation for HeLa S3, overlapped with AcNPM1 peaks. (C) UCSC genome browser snapshot showing AcNPM1 enrichment at TSS and enhancer regions defined by ChromHMM + Segway combined segmentation for HeLa S3 genome. (Key: TSS, predicted promoter region including TSS; E, enhancer). (D) Boxplots showing AcNPM1 read density on AcNPM1 peaks that overlap or do not overlap DNase I hypersensitive sites (DHSs). (E) Boxplots showing AcNPM1 read density on AcNPM1 peaks with high or low enrichment of H3K27ac. (F-G) Boxplots showing AcNPM1 read density on AcNPM1 peaks that overlap or do not overlap (F) p300 and (G) RNA Pol II (Pol2). (H) Transcription factor binding motifs enriched in AcNPM1 peaks and broadly grouped by transcription factor family. *P*-value < 0.05. (I) Percent number of AcNPM1 peaks overlapped with indicated transcription factor and chromatin-associated factor peaks from ENCODE data. (J) Pearson correlation plot of transcription factor and other chromatin-associated factor co-occupancy at AcNPM1 peaks. Hierarchical clustering was performed to identify factors that co-occupy AcNPM1 peaks. Color scale depicts Pearson correlation coefficient values.

### NPM1 functions as a transcriptional coactivator

Given that AcNPM1 co-occupies with RNA pol II as well as TFs at transcriptional regulatory elements, we examined whether NPM1 could interact directly with these factors. Earlier studies have utilized immunoprecipitation (IP)-mass spectrometry approaches from whole-cell lysates to identify NPM1 binding partners. However, the majority of the NPM1 protein in cells is present in the nucleolus, associated with nucleolar proteins as well as ribosomal subunits and proteins. Thus, it might prove difficult to detect the binding partners of the smaller pool of AcNPM1 that interacts with TFs and transcription machinery in the nucleoplasm. We, therefore, used a high-throughput protein-protein interaction profiling approach (Michaud and Snyder 2002) using an array of 9560 human proteins and recombinant NPM1. The interaction was confirmed by detecting NPM1 using a specific antibody and a fluorophore-conjugated secondary antibody. In parallel, a negative control array was probed with only the NPM1 antibody and secondary antibody (see supplemental methods). Using this approach, we identified 3345 proteins that bind with NPM1 significantly (Table S3). We identified several novel binding proteins with roles in transcription as well as other processes (Figure S3A-I, Figure 3A). In order to confirm the validity of these results, we investigated whether known interacting partners of NPM1 were identified in our protein-protein interaction profiling results. Known interacting partners for NPM1 were extracted from the BioGRID database (Stark et al. 2006). Of the 623 known NPM1 interacting partners, only 279 proteins were on the array. Of these, only 143 passed the stringent thresholds we used to detect a positive signal indicating high confidence in the detected NPM1 interacting partners (see supplemental Methods) (Figure S3J). Next, we experimentally validated the results of this approach using four novel candidate proteins identified by the array, namely, transcription factor AP-4, RNA polymerase II subunit K (POLR2K), positive coactivator 4 (PC4), and snail family transcriptional repressor 1 (SNAI1). We generated His_6_-tagged constructs and purified the recombinant proteins from *E. coli*. These His_6_-tagged proteins were used in combination with FLAG-tagged recombinant unacetylated (mock/without p300) or *in vitro* acetylated NPM1 protein in Ni-NTA pull-down assays to determine interaction with AcNPM1 as well. We found that AP-4, POLR2K and PC4 interacted with both unacetylated and acetylated versions of NPM1 (Figure 3B-D) whereas SNAI1 interacted only with unacetylated NPM1 and not AcNPM1 (Figure 3E) in the experimental conditions tested. These results indicate that the interacting proteins identified by this assay can be validated in our *in vitro* interaction assays. We performed gene ontology analyses to identify major biological processes associated with the interacting partners of NPM1. We identified several novel NPM1 interacting proteins that are subunits of different histone acetyltransferase (HAT) complexes (Figure 3F). Moreover, we found RNA Pol II subunits (POLR2C, POLR2E, POLR2I, POLR2K, POLR2M), GTFs (GTF2A2, GTF2E1, GTF2E2, GTF2F1), TFIID associated factors (TAF5L, TAF6, TAF7L, TAF8, TAF9) and Mediator subunits (MED4, MED8, MED11, MED12, MED17, MED19, MED22, MED31) among NPM1 interacting proteins (Figure 3G). These results show that NPM1 interacts with the basal transcriptional machinery and HAT complexes and could potentially function as a coactivator during transcription.

**Figure 3:**
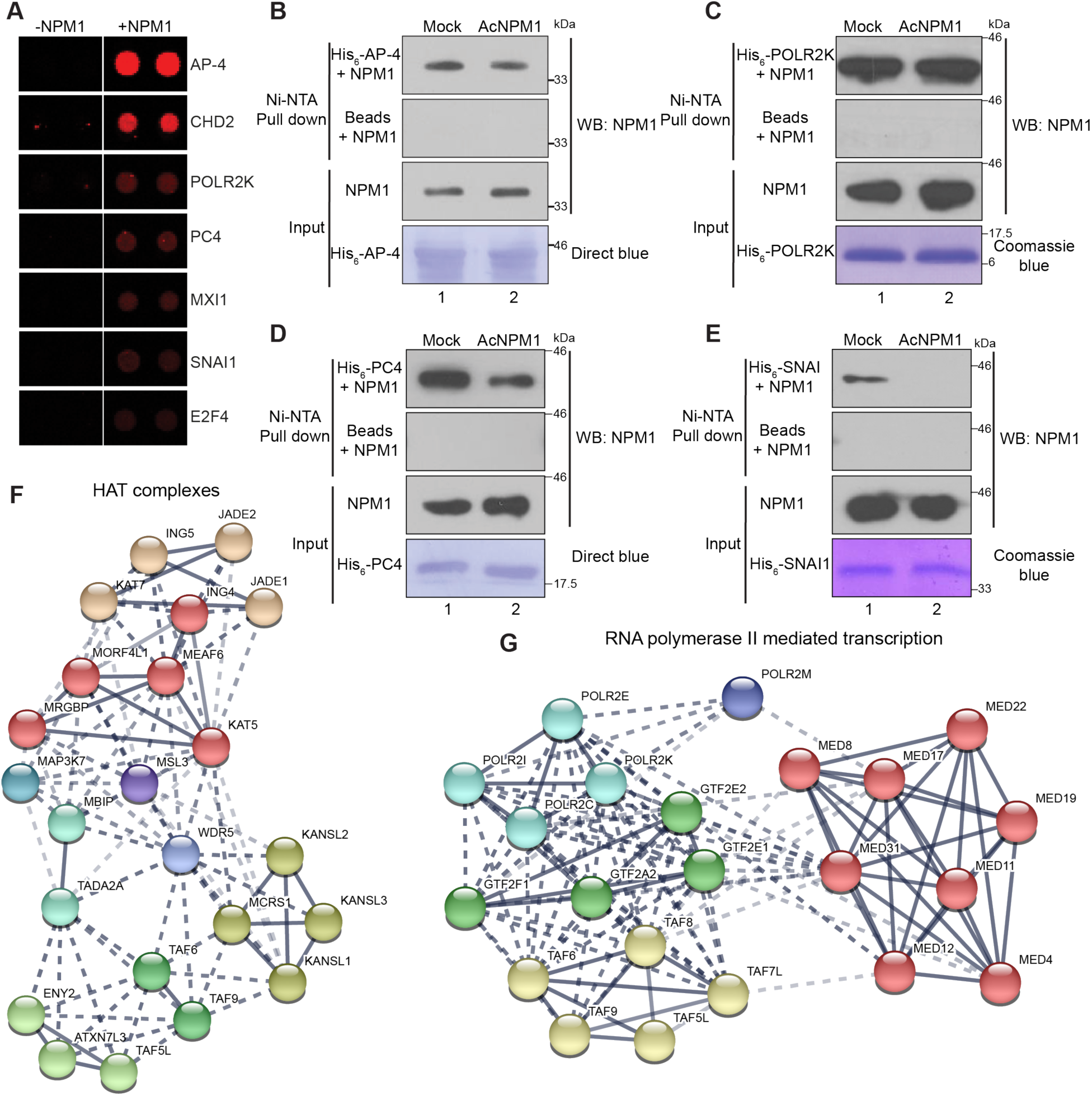
NPM1 functions as a transcriptional coactivator. (A) Representative images of the protoarray probed with (right) or without NPM1 protein (left) showing spots (red) of selected interacting proteins. (B-E) *In vitro* Ni-NTA pull-down of FLAG-tagged recombinant NPM1 (Mock) or acetylated NPM1 (AcNPM1) using (B) His_6_-AP-4 (C) His_6_-POLR2K (D) His_6_-PC4 (E) His_6_-SNAI1 proteins. Western blots were performed with NPM1 antibody after the pull-downs. Equal protein loading is shown using Direct Blue staining of the membrane or Coomassie blue staining of the gel. Input is 20% of NPM1. (F-G) String database (version 11.0) network showing NPM1 interacting proteins identified by protein-protein interaction profiling. Proteins present in HAT complexes are shown in (F) and those involved in RNA Pol II-mediated transcription are shown in (G).

### Histone chaperone activity contributes to transcription regulation by NPM1

In addition to being a transcriptional coactivator, NPM1 is also a histone chaperone and its histone chaperone activity could contribute to transcriptional activation. To test this, we introduced mutations in the N-terminal oligomerization domain of NPM1 that would interfere with the oligomerization property of NPM1 and likely its histone chaperone activity. We targeted highly conserved hydrophobic residues within the N-terminal domain that are effective in abrogating the oligomerization of NPM1 (Figure 4A). NSC348884, a drug that inhibits oligomerization of NPM1 (Qi et al. 2008), binds to residues Tyr17 and Leu18 residues in the NPM1 monomer. Moreover, mutation of Cys21 to Phe (C21F) in NPM1 leads to a loss of the oligomerization property and molecular chaperone activity of NPM1 (Prinos et al. 2011). Tyr17 and Leu18 residues are present in the first β strand of the monomer of NPM1 (Lee et al. 2007) and are necessary for monomer-monomer interaction. Further, these residues have conserved substitutions in other orthologs of NPM1, indicating the functional conservation of these residues for monomer-monomer interactions within the NPM1 pentamer (Figure 4A).

**Figure 4:**
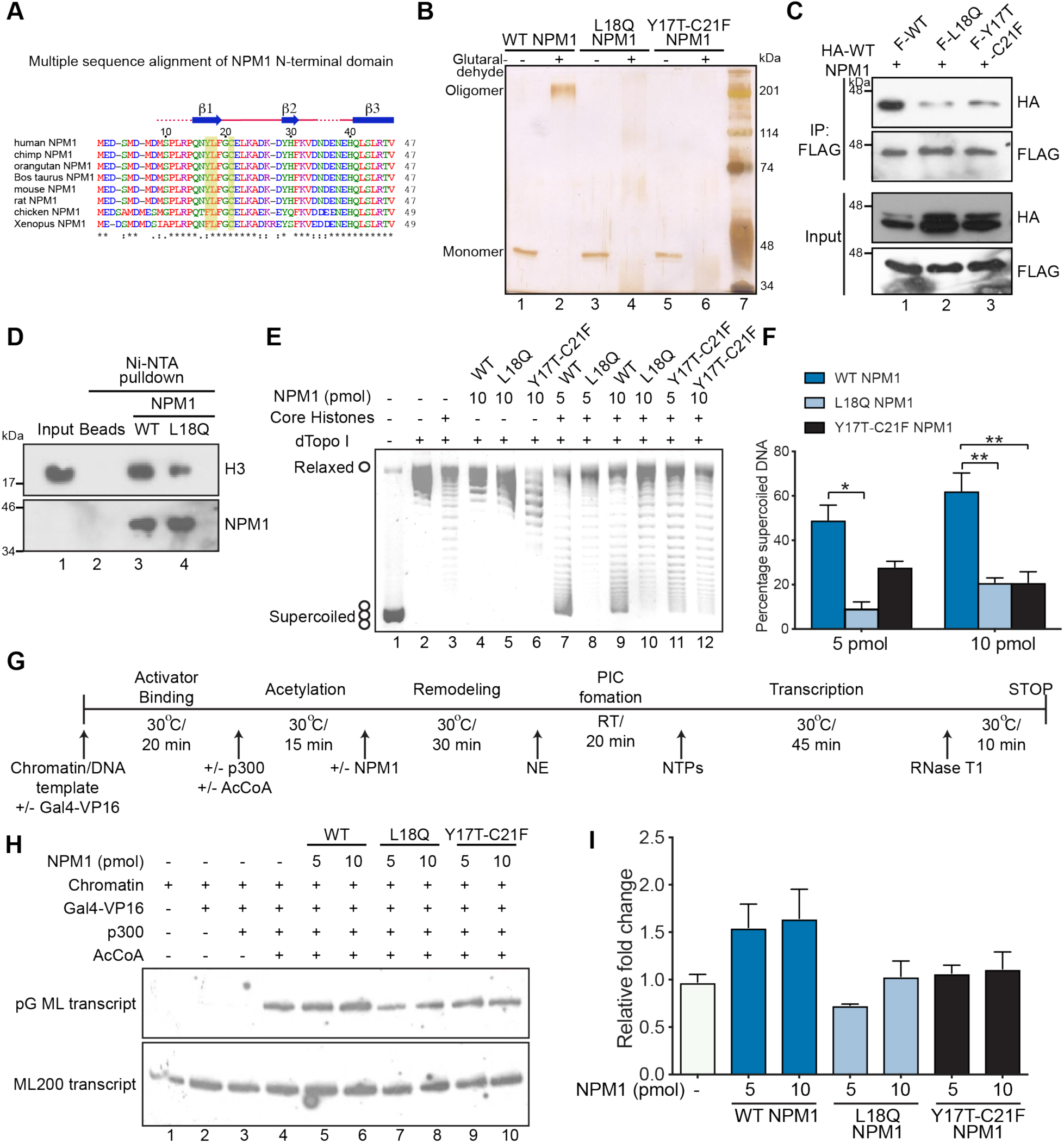
Histone chaperone activity is important for transcription regulation by NPM1. (A) Multiple sequence alignment of the first half of the N-terminal domain of NPM1 from indicated species. Targeted residues are highlighted. The secondary structure of human NPM1 protein is indicated at the top. (B) Silver stained gel showing WT NPM1, L18Q NPM1 and Y17T-C21F NPM1 proteins with or without glutaraldehyde crosslinking. (C) Western blot after anti-FLAG immunoprecipitation in HEK-293 cells co-transfected with HA-tagged WT NPM1 and either FLAG-tagged WT NPM1, L18Q NPM1 or Y17T-C21F NPM1. Top two panels show western blots of immunoprecipitates with HA and FLAG antibodies respectively. Bottom two panels show western blots of Inputs with HA and FLAG antibodies respectively. (D) *In vitro* Ni-NTA pull-down of recombinant histone H3 by His_6_-WT-NPM1 or His_6_-L18Q-NPM1. Ni-NTA beads with H3 serves as negative control. Input is 25% of histone H3 used for the pull-down. (E) Supercoiling assay with 5 and 10 pmol of WT, L18Q or Y17T-C21F NPM1 as indicated. (F) Quantification of the amount of supercoiled DNA obtained after supercoiling assay using WT NPM1, L18Q NPM1, and Y17T-C21F NPM1. The supercoiled DNA formed was quantified using Image J software and expressed as a percentage of the total supercoiled DNA used. Values are Mean + SEM from three independent experiments. Statistical significance was calculated using one-way ANOVA, Tukey’s multiple comparisons test. \**P*-value < 0.05, \*\**P*-value < 0.01. (G) Schematic representation of the *in vitro* chromatin transcription protocol. RT, room temperature; PIC, preinitiation complex; NE, nuclear extract. (H) Chromatin, freshly assembled using the ACF-NAP1 assembly system, was subjected to the protocol described in (G). Lanes 5, 7 and 9 have 5 pmol and lanes 6, 8 and 10 have 10 pmol of WT NPM1 or the respective mutants added as indicated. The relative transcription per lane (in fold activation over the acetylation-dependent transcription lane (lane 4)) was determined by densitometric analysis and presented as a bar graph in (I). Values are Mean + SEM from two independent experiments.

We generated several NPM1 mutant constructs with a combination of mutations in Tyr17, Leu18, and Cys21 (Figure S4A). The Tyr17 was mutated to Threonine (Thr) to remove the aromatic group and Leu18 was mutated to Glutamine (Gln) to replace the hydrophobic side chain with a polar one. These mutants were expressed and purified from *E*. *coli* and tested for their oligomerization ability using a glutaraldehyde cross-linking assay (Figure S4A). The mutant proteins exhibited a range of oligomerization defects, with some showing intermediate forms such as dimers while others aggregated and did not enter the resolving gel (Figure S4A). Of these, two mutants Y17T-C21F and L18Q NPM1 were the most defective in oligomerization as they were present mainly in the monomeric form (Figure S4A, lanes 3 and 4; Figure 4B). To further investigate their oligomerization capability in cells, we generated FLAG-tagged mammalian expression constructs of these mutants. HA-tagged wild type (WT) NPM1 was co-transfected with either FLAG-tagged WT NPM1, L18Q NPM1 or Y17T-C21F NPM1 in HEK-293 cells and co-IPs were performed with FLAG antibody. Results showed that the L18Q NPM1 and Y17T-C21F NPM1 were considerably less efficient in pulling down HA-tagged NPM1 as compared to WT-NPM1 (Figure 4C, compare lanes 2 and 3 versus 1 in top panel). The L18Q mutant showed the least ability to oligomerize with WT NPM1. To rule out the possibility that the mutants are simply less stable than WT NPM1 in cells, we determined the half-life of WT NPM1 protein and compared it to FLAG-tagged versions of Y17T-C21F NPM1 and L18Q NPM1 (Figure S4B-C). We found that half-lives of the mutant proteins are comparable to that of the WT protein (Figure S4B-C). These results indicate that the L18Q and Y17T-C21F mutants are indeed less efficient in forming oligomers as compared to WT NPM1. An in-depth analysis of the crystal structure of NPM1 revealed that the Leu18 residue of one monomer interacts with other hydrophobic residues from the other monomer namely, Ile72, Val74, Leu76, Phe92, Ile94 and are buried together deep within the monomer-monomer interaction interface forming a hydrophobic pocket (Figure S4D). Replacing Leu18 with Gln containing a polar side chain not only makes the side chain of Leu18 unavailable for hydrophobic interactions but also may disturb the hydrophobic pocket. This might explain the observed inability of L18Q NPM1 to form pentamers.

After confirming that the two mutants L18Q NPM1 and Y17T-C21F NPM1 are indeed defective in oligomerization in the experimental conditions tested, we tested their histone interaction and histone chaperone activity. L18Q NPM1 was less efficient in pulling down histone H3 as compared to WT NPM1 (Figure 4D). We next compared the histone chaperone activity of WT NPM1, L18Q NPM1 and Y17T-C21F NPM1 using a histone transfer assay (Senapati et al. 2015) with increasing protein amounts. Both the mutants displayed a reduced ability to deposit histones measured by the extent of supercoiling of plasmid DNA as compared to WT NPM1 (Figure 4E, compare lanes 8, 11 versus 7 and 4F) at both protein concentrations tested (Figure 4E, compare lanes 10, 12 versus 9 and 4F). The results show that oligomerization of NPM1 protein is critical for its histone chaperone activity. We then tested the transcriptional activation property of the histone chaperone-deficient mutants of NPM1 using an acetylation-dependent chromatin transcription assay as indicated in (Figure 4G) (Senapati et al. 2015). The mutant proteins L18Q NPM1 and Y17T-C21F NPM1 were tested in two different concentrations in the transcription assay and compared with WT NPM1. Both the mutant proteins were unable to activate transcription from the chromatin template when compared with the same concentration of WT NPM1 (Figure 4H, compare lanes 7, 9 versus 5, and lanes 8, 10 versus 6, and 4I). Overall these results show that the histone chaperone activity is important for transcriptional activation by NPM1.

### NPM1 is involved in promoting proliferation, migration, and invasion of oral cancer cells

We have earlier reported that NPM1 is overexpressed in OSCC with a concomitant increase in the acetylated NPM1 pool (Shandilya et al. 2009). In order to determine whether the increased AcNPM1 pool is involved in transcriptional regulation of genes in OSCC, we used a shRNA-mediated loss-of-function approach. We examined the expression of NPM1 across various available oral cancer cell lines at mRNA (Figure S5A) and protein (Figure S5B) levels and found high NPM1 expression in the oral cancer cell line AW13516 (Figure S5A-B). To investigate the role of overexpressed NPM1 in oral tumorigenesis, we made an inducible Tet-On NPM1 shRNA cell line in the AW13516 background. Significant downregulation of NPM1 at mRNA (Figure S5C) and protein (Figure S5D) levels was obtained after doxycycline treatment (Dox) with an appreciable decrease in the growth rate of cells as compared to the untreated (UT) cells (Figure 5A). To further investigate the impact of NPM1 knockdown on oral cancer cell proliferation, we performed colony formation assay. NPM1 knockdown did not result in any significant difference in the number of colonies obtained after NPM1 knockdown (Figure 5B). However, the size of the colonies with NPM1 knockdown was severely reduced as compared to the untreated cells, thus indicating that NPM1 might be involved in oral cancer cell proliferation (Figure 5B). To further demonstrate the role of NPM1 in the proliferation of AW13516 cells, we compared wound closure rates by real-time imaging. We found that cells with NPM1 knockdown had slower wound closure rates as compared to the untreated cells (Figure 5C). Wound healing can be attributed to both proliferation as well as migration of the adjacent cells. In order to assess whether NPM1 knockdown might affect the migratory property of oral cancer cells as well, we performed the wound healing assay with mitomycin C pre-treatment to block cell division in cells. We found that NPM1 knockdown cells could not recover the wound whereas the untreated cells partially closed the wound (Figure 5D) post-12 h of wound creation. We further investigated the effect of NPM1 knockdown in the migration and invasion potential of oral cancer cells using transwell assays. We observed a significant reduction in the migration as well as invasion potential of AW13516 cells upon NPM1 knockdown (Figure 5E-F). Overall these results indicate that NPM1 overexpression augments cell proliferation, migration, and invasion during oral tumorigenesis.

**Figure 5:**
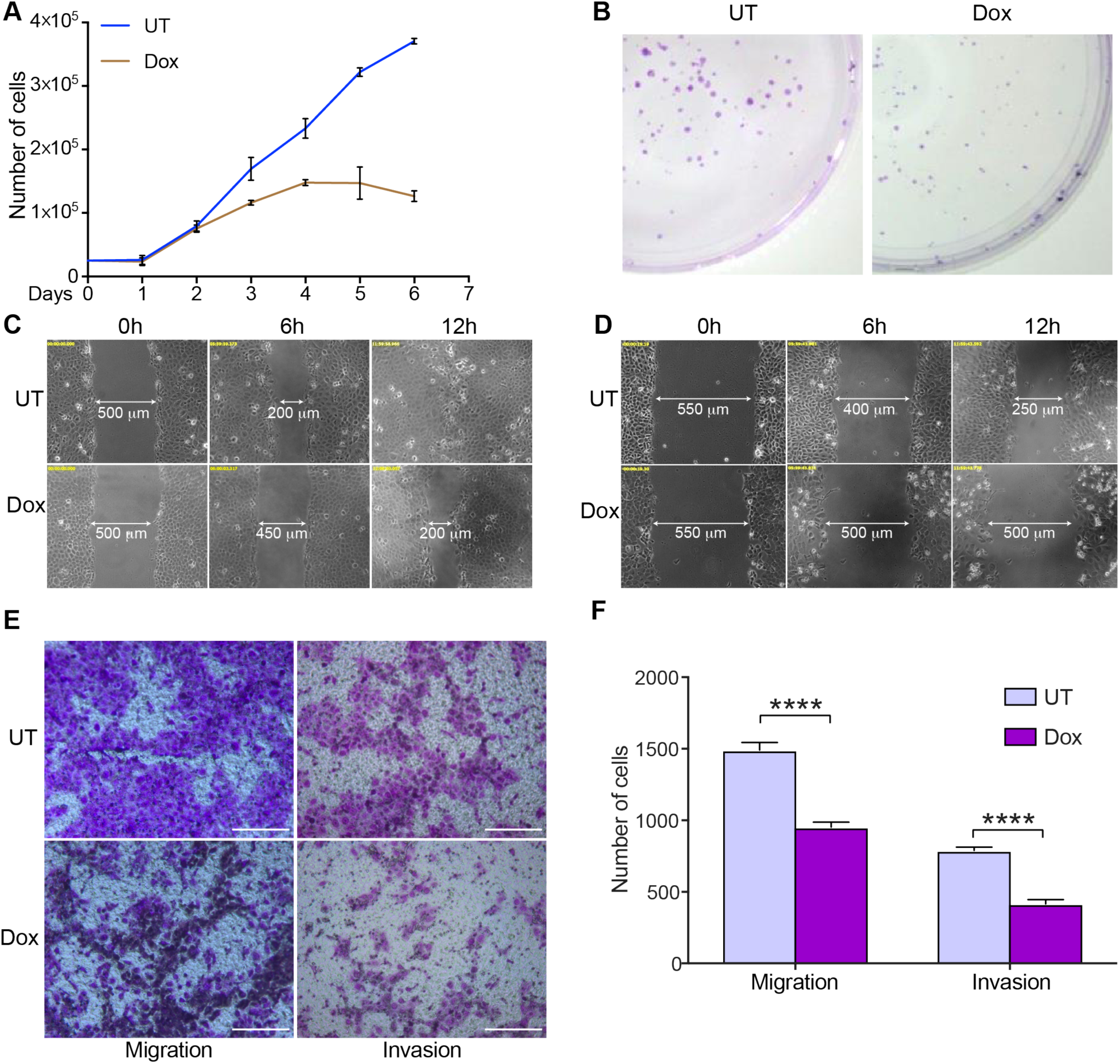
NPM1 is involved in promoting proliferation, migration, and invasion of oral cancer cells. (A) Line graphs represent the number of cells with (Dox) or without doxycycline (UT) induction of NPM1 shRNA in AW13516-shNPM1 cells. Untreated cells were seeded at an initial seeding density was 2.5×10^4^. Values are Mean ± SEM from two independent experiments. (B) Colony formation assay upon doxycycline-induced knockdown of NPM1 (Dox) in AW13516-shNPM1 cells compared to untreated (UT) cells. (C) Photomicrographs of AW13516-shNPM1 cell line either treated (Dox) or untreated (UT) with doxycycline, were captured in real-time for a period of 12 h post-wound creation. Representative images show wound length measured in microns post-6 h and 12 h in untreated (UT) and doxycycline-treated (Dox) conditions. (D) AW13516-shNPM1 cell line was pre-treated with mitomycin C (5 µg/ml) for 2 h followed by wound creation. Representative images show wound length measured in microns post-6 h and 12 h in untreated (UT) and doxycycline-treated (Dox) conditions. (E) Representative images showing migrated or invaded cells from untreated (UT) and doxycycline-treated (Dox) cells in transwell assays. Scale bar is 45 µm. (F) Bar graphs show the quantification of migrated or invaded cells from untreated (UT) and doxycycline-treated (Dox) cells in transwell assays. Values are Mean + SEM from two independent experiments and three fields from each experiment. Statistical significance was calculated using unpaired two-tailed Student’s *t*-test. \*\*\*\**P*-value < 0.0001.

### NPM1/AcNPM1 regulates the gene-network involved in oral tumorigenesis

In order to gain insights into the transcriptional changes brought about by NPM1 knockdown in AW13516 cells, we performed RNA-sequencing with and without doxycycline treatment. NPM1 knockdown leads to significant changes in gene expression (Figure S5E). Of 925 differentially expressed genes, 663 genes were downregulated and 262 were upregulated (Figure 6A). To identify the biological processes affected by NPM1 knockdown, we performed Gene Set Enrichment Analyses (GSEA) (Subramanian et al. 2005). Results showed that genes downregulated upon NPM1 knockdown are enriched in gene sets related to cancer processes namely cell proliferation, negative regulation of apoptosis, cell migration, angiogenesis, hypoxia, inflammation, epithelial-mesenchymal transition (EMT) among others (Figure 6B). Moreover, genes in the AP-1 and HIF-1*α* transcription factor network and genes downstream of TNF-*α* signaling through NF-*κ*B were also enriched (Figure 6B). Interestingly, we also found NF-*κ*B, HIF-1*α* and AP-1 TF binding motifs to be highly enriched in the downregulated gene promoters (Figure S5F). These results indicate that the gene targets of NF-*κ*B, HIF-1*α* and AP-1 TFs are highly downregulated upon NPM1 knockdown (Figure S5G-I) and might be co-regulated by NPM1. Heatmaps of normalized expression of genes show significant downregulation of genes involved In cell proliferation, epithelial-to-mesenchymal transition (EMT), cell migration (Figure 6C), angiogenesis and negative regulation of apoptosis (Figure S5J). We also validated the downregulation of a few candidate genes from these pathways using RT-qPCR analyses (Figure 6D). Significant downregulation of all tested genes including NPM1 was observed after NPM1 knockdown in AW13516 cells. Since AcNPM1 occupies promoter regions and might be involved in the expression of these genes, we tested the AcNPM1 occupancy at the promoters of a few candidate genes with or without NPM1 knockdown (Figure S6K). We indeed observed AcNPM1 occupancy at the promoters of candidate genes downregulated by NPM1 knockdown (Figure 6E-H). Moreover, NPM1 knockdown led to a significant decrease in AcNPM1 occupancy at these gene promoters indicating that AcNPM1 is involved in the transcriptional regulation of these genes (Figure 6E-H).

**Figure 6:**
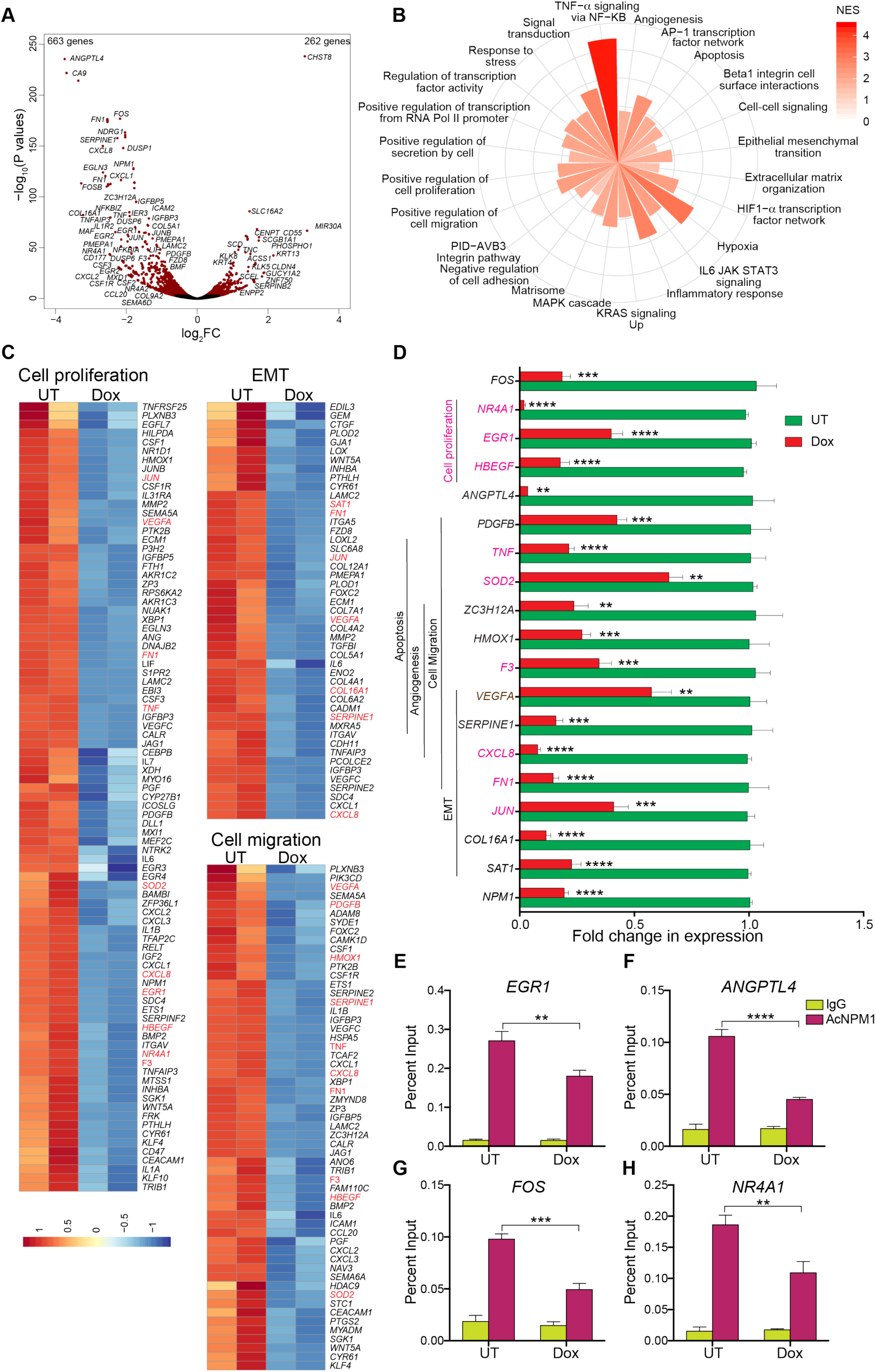
NPM1/AcNPM1 regulates the gene-network involved in oral tumorigenesis. (A) Volcano plot showing genes with significantly altered expression (red) after NPM1 knockdown in AW13516 cells. (B) Gene sets enriched in genes downregulated after NPM1 knockdown. The absolute value of Normalized enrichment score (NES) from Gene Set Enrichment Analysis (GSEA) is shown. *P*-value < 0.05. (C) Heatmaps of normalized read counts at each gene in untreated (UT) and doxycycline-treated (Dox) cells for genes in the “Cell proliferation”, “Epithelial to Mesenchymal transition (EMT) and “Cell Migration” gene sets. (D) Fold change in expression levels of indicated genes as measured by RT-qPCR after NPM1 knockdown by doxycycline treatment (Dox) in AW13516 cells compared to untreated (UT) control. Internal normalization was done with housekeeping gene β-actin levels. Values are Mean + SEM from four independent experiments. Statistical significance was calculated using unpaired two-tailed Student’s *t*-test. \*\**P*-value < 0.01, \*\*\**P*-value < 0.001, \*\*\*\**P*-value < 0.0001. (E-H) AcNPM1 enrichment at indicated gene promoters represented as Percent input and measured by ChIP-qPCR after NPM1 knockdown by doxycycline treatment (Dox) in AW13516 cells compared to untreated (UT) control. Values are Mean + SEM from three independent experiments. Statistical significance was calculated using one-way ANOVA, Tukey’s multiple comparisons test. \*\**P*-value < 0.01, \*\*\**P*-value < 0.001, \*\*\*\**P*-value < 0.0001.

Knockdown of NPM1 affected cell proliferation, migration and invasion in AW13516 oral cancer cell line as well as genes associated with these processes. We next investigated whether NPM1 knockdown would affect tumor growth in orthotopic tumor models in mice. We generated a stable cell line expressing the luciferase gene in the AW13516-shNPM1 background to monitor tumor growth using live imaging in mice. The AW13516-shNPM1-luc+ cells were injected into the floor of the mouth region of the mice, and tumors were allowed to grow for 5 days. One group of mice was then fed with doxycycline (Dox) to induce NPM1 shRNA expression whereas the other group was fed with the vehicle (Veh). Tumor flux (luciferase activity) was measured for the mice from each group using a live animal imaging system. We observed a significant decrease in the tumor flux at day 16 or 11 days after doxycycline treatment (Figure 7A-B) indicating that NPM1 knockdown indeed results in slow growth of tumors. To rule out any adverse effects of doxycycline on oral tumors, we performed a parallel experiment using a control group of mice injected with AW13516-luc+ cells (lacking the NPM1 shRNA) and treated as described above. We did not find any significant difference in the tumor flux among the vehicle and doxycycline fed animals in this group (Figure S6A-B), indicating no effect of doxycycline alone on oral tumor growth. We next tested the levels of NPM1 and Ki67, a proliferation marker, in the tumors from the Veh and Dox groups. As expected, we observed a decreased staining of NPM1 (Figure 7C-D) as well as the proliferation marker Ki67 (Figure 7C) in tumors from the Dox group as compared to the Veh group. Since we observed a decrease in EMT genes in the RNA-seq data after NPM1 knockdown, we also stained the orthotopic tumors with E-cadherin and Fibronectin. E-cadherin, an epithelial marker, increased in contrast to the mesenchymal marker Fibronectin, whose levels decreased upon NPM1 knockdown showing inhibition of EMT (Figure 7E). This is consistent with reduced invasive property of cells after NPM1 knockdown in transwell assays. In support of this finding, we did notice that some of the animals in the Veh group showed adjoining lymph node metastasis which was appreciably less in the Dox group (Figure S6C). To investigate whether knockdown of NPM1 has any effect on the oral cancer stem cell population, we stained the tumors from the Veh and Dox groups with CD44, a stem cell marker. Tumor samples from Dox group showed an evident reduction in CD44-positive cells (Figure 7F) indicating a loss of oral cancer stem cell population. In summary, these results suggest that NPM1 overexpression in OSCC might potentiate the expression of tumor-promoting genes through AcNPM1-mediated transcriptional regulation.

**Figure 7:**
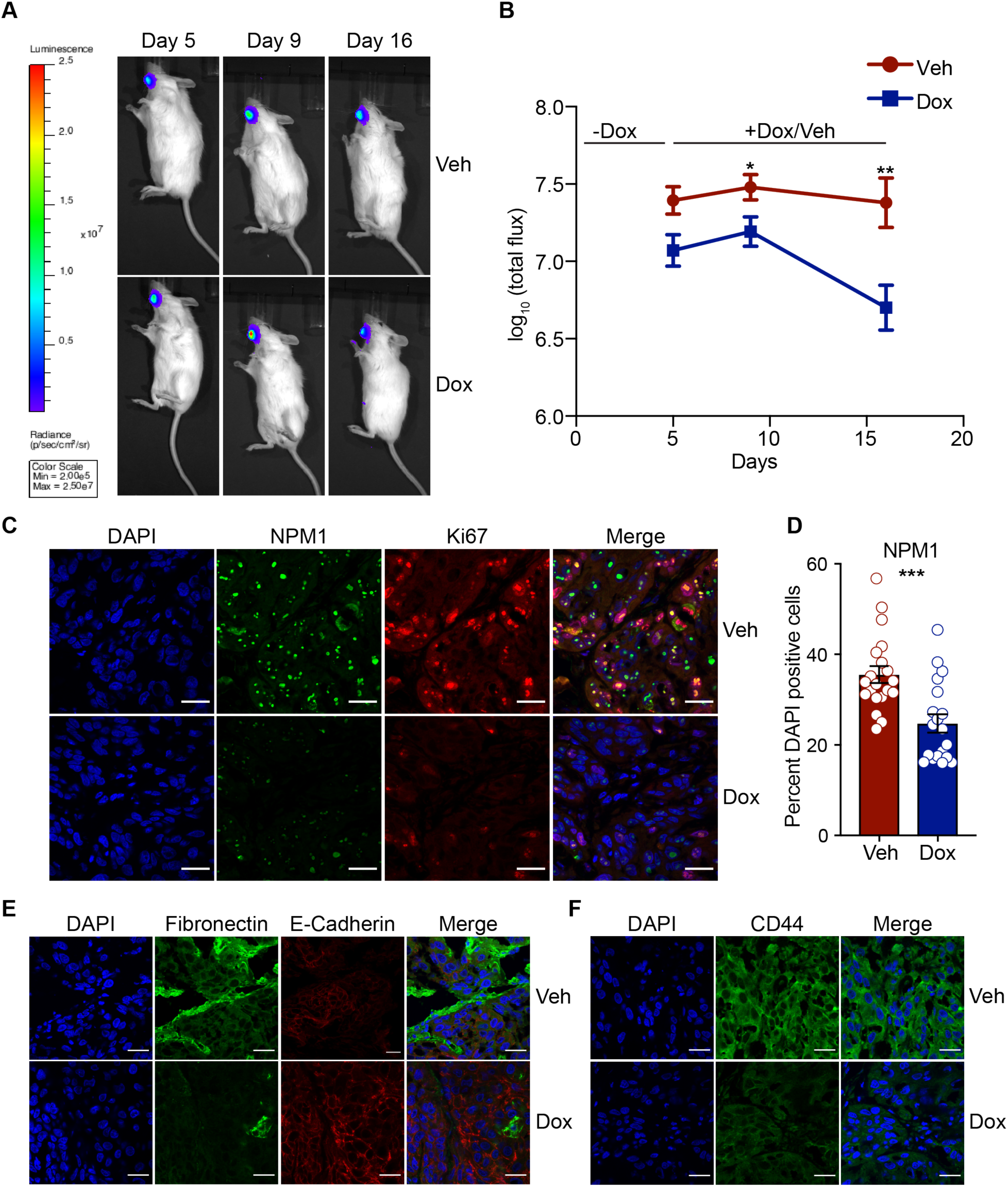
NPM1 knockdown abrogates oral tumor growth in mice. (A) Representative bioluminescent imaging of Vehicle (Veh) or doxycycline fed (Dox) mice at 5, 9 and 16 days post-injection with 1*×*10^6^ AW13516-shNPM1-luc+ cells. Units represent relative light units. (B) Bioluminescence intensity measured at 5, 9 and 16 days post-injection of the cells, for the Veh and Dox groups are shown in the graph. Values are Mean ± SEM of log_10_(total flux), n = 7 animals in each group. Statistical significance was calculated using unpaired two-tailed Student’s *t*-test. \**P*-value < 0.05, \*\**P*-value < 0.01. (C) Representative images showing immunohistochemical staining of tumor tissues from Veh and Dox groups with NPM1 and Ki67 antibodies. (D) Quantification of NPM1-positive cells expressed as a percentage of DAPI-positive cells. N = 5 animals from each group with 3-4 fields from each animal. Statistical significance was calculated using two-tailed Mann-Whitney test. \*\*\**P*-value < 0.001. (E) Representative images showing immunohistochemical staining of tumor tissues from Veh and Dox groups with Fibronectin, E-Cadherin, and (F) CD44 antibodies. (C, E, and F) Nuclei are stained with DAPI. Magnification is 10X and scale bar is 25 µm.

## DISCUSSION

Histone chaperones are emerging as new players in regulating histone dynamics during transcription. The role of histone chaperone, FACT (facilitates chromatin transcription) in the process of nucleosome assembly and disassembly during transcription initiation and elongation is well documented (Formosa 2012). NPM1, a member of the Nucleoplasmin family (Lee et al. 2007), was identified to be a histone chaperone and a transcriptional coactivator (Okuwaki et al. 2001; Swaminathan et al. 2005). Since then, very few details have emerged regarding the mechanism of NPM1-mediated transcription regulation. NPM1 is acetylated by p300/KAT3B and acetylation enhances its histone chaperone and transcription activation property (Swaminathan et al. 2005). The crystal structure of NPM1, and other proteins from the histone chaperone family, Nucleoplasmin and NO38 were solved, and this has helped in understanding how NPM1 might bind to histones (Dutta et al. 2001; Namboodiri et al. 2004; Lee et al. 2007; Platonova et al. 2011). In this study, we have investigated the mechanisms by which NPM1 might regulate transcription apart from and including its histone chaperone activity. Additionally, we have undertaken a genome-wide approach to identify the transcriptional targets of AcNPM1. ChIP-seq analyses revealed that AcNPM1 occupies transcriptional regulatory elements including promoters and enhancers. AcNPM1 peaks showed a good correlation with histone marks associated with transcriptional activation such as H3K9ac, H3K27ac, and H3K4me3. Interestingly, we also found AcNPM1 peaks on enhancer regions showing high occupancy of H3K27ac, p300, and RNA Pol II, indicating that AcNPM1 is associated with active transcription at promoters and enhancers.

It is well understood that transcription through chromatin *in vivo* is assisted by ATP-dependent remodeling enzymes, histone-modifying enzymes and histone chaperones (Li et al. 2007). Some factors such as RSC (Chromatin structure remodeling) interact with the RNA Pol II subunit Rpb5 (Soutourina et al. 2006) and occupy promoters and ORFs (open reading frames) of genes (Ng et al. 2002). The remodeler SWI/SNF travels with RNA Pol II *in vivo*, evicting histones on active genes while it is transcribing (Schwabish and Struhl 2007). Similarly, histone chaperones Asf1 (Schwabish and Struhl 2006), Spt6 (Kaplan et al. 2003), and FACT (Mason and Struhl 2003), also travel with RNA Pol II during transcript elongation and assist in histone removal and re-deposition after Pol II. We found AcNPM1 to be primarily enriched at promoter regions of genes and not on gene bodies indicating a role in transcription initiation. We cannot rule out the possibility, however, that unmodified NPM1 may be present on gene bodies as multiple attempts to perform NPM1 ChIP-seq failed due to the unavailability of ChIP-grade polyclonal antibodies against NPM1.

NPM1 may also be involved in the recruitment of GTFs or the transcription machinery for initiation of transcription. Indeed, we found that NPM1 interacts with some GTFs (GTF2A2, GTF2E1, GTF2E2, GTF2F1), TFIID associated factors (TAFs), RNA Pol II subunits and Mediator subunits. The TFIID complex binds to the TATA box through TBP and is responsible for the proper positioning of the pre-initiation complex (Burley and Roeder 1996). At TATA-less promoters, the subunits of TFIID bind to various TFs that help the positioning of TFIID (Grunberg and Hahn 2013). TFIIF is recruited along with RNA Polymerase II during initiation of transcription (Lee and Young 2000). Using ChIP-seq data from ENCODE consortium in HeLa S3 cells, we find significant overlap (>40% AcNPM1 peaks) between AcNPM1 peaks and TBP, TAFII-250, two subunits of the TFIID complex (Figure 2I). Similarly, we found an appreciable overlap of AcNPM1 with TFIIF-alpha (GTF2F1) peaks (Figure 2I). Interestingly, 90% of genes downregulated after NPM1 knockdown in oral cancer cells contain a TATA motif on their promoters (Figure S5F). These results provide further confirmation for the protein-protein interaction profiling results and suggest that AcNPM1 might interact with these GTFs to recruit RNA Pol II during initiation (Figure S7). We indeed find ∼52% of AcNPM1 peaks to be co-occupied by RNA Pol II (Figure 2I).

We did not find enrichment of AcNPM1 on rDNA regions (data not shown), unlike unmodified NPM1, which was shown to bind rDNA regions (Murano et al. 2008). This is consistent with the fact that the majority of the NPM1 pool is present in the nucleolus whereas the AcNPM1 pool is localized in the nucleoplasm and may be associated with active transcription machinery (Shandilya et al. 2009) (Figure S7). Similar observations were reported for another nucleolar protein Nucleolin that also undergoes acetylation. ChIP-seq studies using Nucleolin antibody showed its enrichment at the rDNA regions (Cong et al. 2012) whereas acetylated Nucleolin (AcNCL) ChIP-seq revealed that AcNCL was not enriched on rDNA. Rather, AcNCL was localized in nuclear speckles and co-localized with the splicing factor SC35, indicating its probable role in transcription or post-transcriptional processing (Das et al. 2013).

We further demonstrated that NPM1 histone chaperone activity contributes to its transcriptional activation ability. Given that AcNPM1 exhibits higher nucleosome disassembly activity (Swaminathan et al. 2005) and localizes with RNA Pol II at TSS regions, it is possible that AcNPM1 helps in nucleosome disassembly during transcription initiation apart from the recruitment of GTFs and RNA Pol II. Moreover, we found a significant overlap of AcNPM1 peaks with chromodomain helicase DNA-binding (CHD2), an ATP-dependent chromatin remodeler (Liu et al. 2015), from our ChIP-seq data (Figure 2I). CHD2 protein also interacted strongly with recombinant NPM1 in our protein-protein interaction profiling assay (Figure 3A). These results indicate that AcNPM1 might associate with chromatin remodelers to promote nucleosome disassembly (Figure S7).

We found that NPM1 regulates several genes related to cell proliferation, migration, and invasion potential of oral cancer cells. Knockdown of NPM1 decreases the occupancy of AcNPM1 on promoters of these genes, as well as leads to a decrease in their expression (Figure S7). Levels of NPM1 and AcNPM1 are higher in OSCC tissues than adjacent normal tissues (Shandilya et al. 2009). However, the role of the increase in NPM1 and AcNPM1 levels was not well understood. NPM1 overexpression may drive rDNA transcription and ribosome biogenesis. Further, overexpressed NPM1 might act as a molecular chaperone and cater to the increased protein synthesis needs in the rapidly proliferating cancer cells, thereby preventing a misfolded protein stress response. Additionally, the highly acetylated pool of NPM1 in the nucleoplasm drives the expression of genes involved in tumorigenesis. Presumably, NPM1 contributes to tumorigenesis through these mechanisms.

Our protein-protein interaction profiling approach identified several centromeric proteins as novel binding partners of NPM1 (Figure S3I). NPM1 was earlier reported to be associated with the CENP-A nucleosomal complex (Foltz et al. 2006). Previously, we reported that NPM1 relieves the repression of CENP-A assembled chromatin in *in vitro* transcription assays (Shandilya et al. 2014). Hence, NPM1 might be an important factor that regulates centromeric chromatin. Thus, our protein-protein interaction profiling results greatly expands our knowledge of the NPM1 interactome. The significance of this is due to the fact that NPM1 protein exhibits intrinsically disordered regions (IDRs) that enable its role as a chaperone and multifunctional protein. We identified several novel binding partners of NPM1 in diverse physiological processes where NPM1 is involved, for example, DNA replication, transcription-coupled nucleotide excision repair, cell cycle (Figure S3) to name a few. These results open up new avenues for studying NPM1 function in these processes. Moreover, NPM1 is a nucleolar protein that can undergo liquid-liquid phase separation through multiple mechanisms (Mitrea et al. 2018). A recent study has shown how this property is involved in the ribosomal biogenesis function of NPM1 (Mitrea et al. 2018). Emerging research in the transcription field suggests that the process of transcription occurs in phase-separated condensates where the IDRs of coactivators such as BRD4, MED1 as well as C-terminal domain of RNA Pol II contribute to the phase separation (Cho et al. 2018). Similar to the role of NPM1 in facilitating ribosome biogenesis in the nucleolus, we can surmise that NPM1 could function in forming these phase-separated condensates during transcription through its interactions with RNA as well as a multitude of other coactivators and TFs. However, this will need further experimental validation. Overall, our study provides evidence for the role of histone chaperone NPM1 in transcriptional regulation, advances the knowledge of NPM1 interactome and paves the way for further exploration into these areas.

## METHODS

### Cell Culture

Human cell lines, HEK-293 (CRL-1573) and HeLa S3 (CCL-2.2) were purchased from American Type Culture Collection (ATCC, Manassas, VA, USA) and were grown in Dulbecco’s modified Eagle’s medium (DMEM) (Sigma, St. Louis, MO, USA) at 37°C, 5% CO_2_ in a humidified chamber. UPCI:SCC40 and UPCI:SCC122 oral cancer cell lines were kind gifts from Prof. Susanne M. Gollin (University of Pittsburgh, USA). Oral cancer cell lines AW13516, AW8507, OT9, NT8e, and the esophagus normal Het-1A cells were kind gifts from Dr. Amit Dutt (ACTREC, Mumbai, India). All oral cancer cell lines were grown in Minimum Essential Medium (MEM) (Gibco, Carlsbad, CA) supplemented with 1X non-essential amino acids (Sigma). Het-1A cells were grown in Bronchial Epithelial Cell Growth Medium (BEGM) (Lonza/Clonetics Corporation, Basel, Switzerland). All the media except BEGM were supplemented with 10% Fetal Bovine Serum (v/v) (Life Technologies, Bengaluru, India) and 1X antibiotics containing penicillin, streptomycin, and amphotericin (HiMedia, Mumbai, India). All cell lines used in this study were routinely tested for *Mycoplasma* contamination and used for no more than 10 passages.

NPM1 knockdown cell line was generated in AW13516 cell line by transfecting with NPM1 shRNA construct in pTRIPZ vector (Thermofisher Scientific, Waltham, MA, USA) for a period of 24 h followed by antibiotic selection using 1.6 µg/ml puromycin for 4 days. Colonies were picked and grown under antibiotic selection. The stable cell lines were then characterized for expression of shRNA by performing doxycycline induction at 2 µg/ml and scoring for the expression of TurboRFP in a fluorescence microscope. Clones with high expression of shRNA after doxycycline induction were expanded and sorted using a BD FACSAria™ III (BD Biosciences-US, Franklin Lakes, NJ, USA) cell sorter to select for the high TurboRFP expressing cells. These cells were maintained in MEM complete medium supplemented with L-glutamine and 1.1 µg/ml puromycin.

pGL4 luc neo construct (Promega, Madison, WI, USA) was transfected into the AW13516-shNPM1 cell line for a period of 24 h followed by antibiotic (G418) selection using 800 μg/ml for 7 days. Colonies were picked and grown under antibiotic selection. The stable cell lines were then characterized for expression of luciferase by performing a luciferase assay. Clones with high luciferase activity were expanded, shRNA expression was induced by doxycycline (2 μg/ml) and sorted using a BD FACSAria™ III (BD Biosciences-US) cell sorter to select for the high TurboRFP expressing cells. The cells were maintained in MEM complete medium supplemented with L-glutamine, 800 μg/mL of G418 and 1.1 μg/ml of puromycin.

### Generation of polyclonal antibodies against acetylated NPM1 (K229, K230)ac

A peptide containing acetylated Lys229 (K229) and Lys230 (K230) residues was designed and conjugated with Keyhole Limpet Hemocyanin (KLH) (Genemed Synthesis Inc., San Antonio, TX, USA) KLH-C-KGQESFK(Ac)K(Ac)QEKTP (residues 223 to 235). Polyclonal antibody against the peptide was raised in New Zealand white rabbit following the standard regime for priming and booster doses for antibody generation. The serum was further purified by peptide-affinity chromatography and the specificity of the antibody was confirmed using western blotting, immunofluorescence and dot blot assays.

### Chromatin Immunoprecipitation (ChIP)

ChIP was performed as described earlier (Senapati et al. 2018). Briefly, 10-15 million HeLa S3 or AW13516-shNPM1-luc+ cells were cross-linked using 1% formaldehyde followed by cell lysis in SDS lysis buffer (1% SDS, 10 mM EDTA, 50 mM Tris-HCl, pH 8). Lysates were sonicated in a Diagenode Bioruptor (Liège, Belgium) to produce 100–300 bp DNA fragments and precleared prior to immuno-precipitation with specific antibodies and pre-blocked Protein G-Sepharose (Amersham Biosciences, Buckinghamshire, UK) for 1 h. Precleared cell lysates were incubated with 5 μg of purified AcNPM1 anybody or pre-immune IgG per sample, and pre-blocked protein G-Sepharose beads overnight at 4°C. Beads were then washed successively with low-salt buffer (0.1% SDS, 1% Triton X-100, 2 mM EDTA, 20 mM Tris–HCl, pH 8, and 150 mM NaCl), high-salt buffer (0.1% SDS, 1% Triton X-100, 2 mM EDTA, 20 mM Tris–HCl, pH 8, and 500 mM NaCl), LiCl buffer (250 mM LiCl, 1% NP40, 1% NaDOC, 1 mM EDTA, and 10 mM Tris–HCl, pH 8), and TE (10 mM Tris–HCl, pH 8, and 1 mM EDTA). DNA-protein complexes were recovered from beads in elution buffer (0.1% SDS and 100 mM NaHCO_3_). The ChIP eluates and input samples in the elution buffer were then reverse cross-linked by adding 200 mM NaCl, 20 µg Proteinase K (Sigma), at 65°C for 4 h. Subsequently, 20 µg of RNase A (Sigma) were added and the samples were further incubated for 15 min at 37°C. The immunoprecipitated DNA was extracted using phenol-chloroform, ethanol precipitated and used for quantitative PCR. For sequencing, the extracted ChIP DNA was further re-purified using NucleoSpin® Gel and PCR Clean-up kit (Macherey-Nagel, Düren, Germany). The region-specific primer sets used for the ChIP-qPCR analysis have been mentioned in Table S2.

### ChIP-seq analyses

Library preparation and sequencing were performed at GATC Biotech (Konstanz, Germany) on a HiSeq 2500 Illumina sequencer. Sequencing reads from each library were adapter trimmed using Trim Galore (https://github.com/FelixKrueger/TrimGalore) and then aligned to the human hg19 assembly using bowtie2 (Langmead and Salzberg 2012) using parameters --phred33 --local –N 1. Alignment rates of more than 89% were obtained. The alignment statistics are reported in Table S1. Peak calling was performed using MACS2/2.1.1.20160309 (Zhang et al. 2008) using --broad option with default parameters. We identified 24660 AcNPM1 peaks conserved between the two replicates after filtering out the hg19 blacklisted regions (Amemiya et al. 2019). Peaks were assigned to nearest RefSeq TSS using annotatePeaks.pl from homer package (Heinz et al. 2010). Random peaks were obtained by using shuffle from the BEDTools suite (Quinlan and Hall 2010). Bigwig files were generated for visualization using deepTools (Ramirez et al. 2014). Aggregate plots and heatmaps were generated using deepTools. Broadpeak and bigwig files for ENCODE histone modification ChIP-seq data for HeLa S3 cells were downloaded from GEO (GSE29611). Jaccard indexes were calculated using jaccard function in BEDtools. Combined segmentation bed file for HeLa S3 was downloaded from UCSC genome browser (http://hgdownload.cse.ucsc.edu/goldenpath/hg19/encodeDCC/wgEncodeAwgSegmentation/wgEncodeAwgSegmentationCombinedHelas3.bed). Motif enrichment analyses were performed using homer. BED files for DNase I hypersensitivity sites for HeLa S3 were downloaded from UCSC genome browser (http://hgdownload.cse.ucsc.edu/goldenPath/hg19/encodeDCC/wgEncodeAwgDnaseUniform/wgEncodeAwgDnaseUwdukeHelas3UniPk.narrowPeak.gz). BED files for transcription factor ChIP-seq data were downloaded from GEO (GSE33213, GSE32465, GSE3147).

### RNA-seq analysis and reverse transcription (RT)-qPCR

Total RNA was extracted from untreated (UT) or doxycycline-treated (Dox) (1 µg/ml for 6 days) AW13516-shNPM1-luc+ cells using the GenElute Mammalian Total RNA Miniprep Kit (Sigma) followed by On-Column DNase I Digestion as per manufacturer’s protocol. RNA integrity was measured by Agilent Bioanalyzer 2100 (Agilent Technologies, Santa Clara, CA, USA). RNA-seq libraries were prepared using the KAPA Stranded mRNA-Seq Kit (KAPA Biosystems, Wilmington, MA, USA, Cat. No. KK8421) from 250 ng of total RNA. Library quality was assessed on an Agilent Bioanalyzer 2100 and quantified using a Qubit 3.0 Fluorometer (Life Technologies). Sequencing was performed using an Illumina HiSeq 4000 (Illumina Inc., San Diego, CA, USA) at Quick Biology Inc. (Pasadena, CA, USA; via Science Exchange, Palo Alto, CA, USA) in paired-end mode. About 24-32 million paired-end reads of length 150 were obtained from each library (Table S1). Raw sequences were aligned to the hg19 reference genome using HISAT2 2.1.0 (Kim et al. 2015) using default parameters. Stringtie 1.3.4 (Pertea et al. 2015) was used with default parameters to assemble transcripts using the RefSeq annotation. Assembled transcripts from all libraries were further merged using --merge option in Stringtie. Merged transcript abundances were measured using bedtools coverage and DESeq2 package (Love et al. 2014) was used to normalize counts and identify differentially expressed genes (log2 fold change *≥* 0.5 and padj < 0.1). Gene Set Enrichment Analysis (GSEA) (Subramanian et al. 2005) was used to determine significantly altered gene ontology and pathways.

For RT-qPCR assays, total RNA was extracted from treated cells using TRIzol reagent (Invitrogen, Carlsbad, CA, USA) according to the manufacturer’s instructions. RNA was treated with DNase I (NEB, Ipswich, MA, USA) according to the manufacturer’s instructions followed by re-precipitation. First-strand cDNA was synthesized from 2 µg of total RNA using Moloney Murine Leukemia Virus Reverse Transcriptase (Sigma) and oligo dT (Sigma) as per the manufacturer’s instructions. Real-time PCR was performed on a Step-One Plus Real-Time PCR Detection System (Applied Biosystems, Waltham, MA, USA) using 2X Power SYBR Green Mastermix (ABI) and the respective specific primers enlisted in Table S2. The data were analyzed using StepOne Software version 2.3. Fold changes were calculated using the formula 2^-(Ct_treated_-Ct_control_). β-actin (*ACTB*) or 18S rRNA were used as housekeeping genes.

### Orthotopic mouse tumor experiments

All animal experiments were performed under the approval of Institute Animal Ethics Committee (IAEC) of RGCB (IAEC/271/TM/2015). AW13516-shNPM1-luc+ cells were orthotopically injected into 2-month old male NOD.CB17-*Prkdc^scid^*/J mice (n=14). A control group of SCID mice (n=5) were injected with AW13516-luc+ cells. Briefly, 1*×*10^6^ cells were resuspended in MEM containing Matrigel Matrix Growth Factor Reduced, Phenol Red-Free (BD Biosciences, Cat No. 356231) and then injected into the floor of the mouth region superficial to the mylohyoid muscle of the anesthetized mice. The AW13516-shNPM1-luc+ injected mice were divided into two groups of seven animals each. For one of these groups, enteral administration of doxycycline (1 mg/ml) was initiated from the fifth day after injection, by dissolving doxycycline in 5% sucrose solution. The solution was changed every 24 h. The control group (Veh) was given only 5% sucrose solution (vehicle). For the negative control, the AW13516-luc+ injected mice were divided into two groups, one having two (provided with the vehicle sucrose solution) and the other having three animals (provided with doxycycline containing sucrose solution).

### Bioluminescence imaging

The animals were anesthetized with 3% isofluorane, intraperitoneally injected with firefly D-luciferin (Promega, Cat No. P1042) (2 mg/150 µl of sterile saline solution) and imaged using IVIS SPECTRUM *in vivo* imaging system (Perkin Elmer, Waltham, MA, USA). For bioluminescence imaging, the acquisition time was set as 4 s for firefly luciferase. Signal intensities were calculated with IVIS SPECTRUM imaging software Living Image, Version 4.3.1 and expressed as photons per second per cm^2^ per steradian. Bioimaging was performed on the fifth, ninth and sixteenth day after injection. For testing lymph node positivity as a sign of cancer metastasis, 3-dimensional bioimaging was performed by dark lock-in thermography (DLIT).

### Immunohistochemistry

Tumors from mice were stored in 4% paraformaldehyde for 24 h after which they were cryoprotected in 30% sucrose solution for two weeks at 4 °C. Cryosections were performed at 7 µm sections using Cryostat LeicaCM1850 UV (Leica Biosystems, Wetzlar, Germany). For staining, the tissue sections were washed with PBS followed by antigen retrieval with 0.01 M citrate buffer (pH 6). The tissues were then permeablized with 0.3% Triton X-100/PBS (PBST) and blocked with 2% serum followed by incubation with primary antibodies overnight at 4°C. Staining was performed in combinations of NPM1 (mouse monoclonal, in-house generated) and Ki67 (Abcam, Cambridge, UK, Cat. No. ab15580), E-cadherin (Santacruz Biotechnology Inc., Dallas, TX, USA, Cat. No. sc-7870) and Fibronectin (Santacruz Biotechnology Inc., sc-271098). CD44 (BD Biosciences, Cat No. 555476) staining was performed separately. The next day, secondary antibody incubation was carried out for 1 h at RT followed by staining of the nuclei with DAPI (Sigma, Cat. No. 32670, Lot No. BCBD9022V) and mounting of the sections with ProLong Gold Antifade Mountant (Invitrogen, Cat. No. P36930, Lot No. 1346597). Images were acquired using NikonA1R LSCM confocal microscope (Nikon, Tokyo, Japan). For NPM1 level quantitation, the total number of nuclei in each field was calculated using Image J software, followed by counting the number of nuclei which were positively stained for NPM1.

### Statistical analyses

Statistical analyses were performed using GraphPad Prism 8.0.2 software (GraphPad Prism Software Inc., San Diego, CA, USA) and R. Shapiro-Wilk normality test was used to confirm the normal distribution of data. For normally distributed data, unpaired two-tailed Student’s *t*-test was used for comparing two means, and one-way analysis of variance (ANOVA) followed by Tukey’s post-hoc test was used for comparison between three or more groups. In case of *t*-tests, equal variance in the groups compared was confirmed using *F* test. For data with significantly different variances, a *t*-test with Welch correction was performed. For non-normally distributed data, Mann–Whitney test was performed for comparison between two groups and Kruskal–Wallis test followed by Dunn’s multiple comparisons test was performed for comparing more than two groups. A *P*-value of < 0.05 was considered statistically significant. Figures were generated using Adobe Illustrator software (Adobe Inc., San Jose, CA, USA).

## DATA ACCESS

All raw and processed sequencing data generated in this study have been submitted to the NCBI Gene Expression Omnibus (GEO; http://www.ncbi.nlm.nih.gov/geo/) under accession number GSE132849.

## ACKNOWLEDGMENTS

This work was supported by Jawaharlal Nehru Center for Advanced Scientific Research (JNCASR), Sir JC Bose Fellowship, Department of Science and Technology (DST), India (Grant No. SR/S2/JCB-28/2010), the Department of Biotechnology (DBT), India (Programme Support on ‘Chromatin and Disease’, Grant No. BT/01/CEIB/10/III/01 and Virtual National Oral Cancer Institute, Grant No. BT/PR17576/MED/30/1690/2016). PS is supported by the Council of Scientific and Industrial Research (CSIR) and, SD1 and AB are supported by the University Grants Commission (UGC), India respectively. We are grateful to Prof. Susanne M. Gollin (University of Pittsburgh, USA) for providing the UPCI:SCC40 and UPCI:SCC122 cell lines and Dr. Amit Dutt (ACTREC, Mumbai, India) for providing the AW13516, AW8507, OT9, NT8e, and the esophageal normal Het-1A cell lines. We also acknowledge the Confocal Imaging Facility and the Flow Cytometry facility at JNCASR, Bangalore, India.

## AUTHOR CONTRIBUTIONS

PS designed and performed most of the experiments, analyzed and interpreted the ChIP-seq, RNA-seq and protein-protein microarray data and wrote the manuscript. SD1 generated the AcNPM1 antibody, performed dot blot assays, RT-qPCR experiments, ChIP-qPCR experiments and analyzed the data. DS performed site-directed mutagenesis experiments, prepared recombinant proteins, performed glutaraldehyde crosslinking experiments and *in vitro* Ni-NTA pull-down experiments. AB performed the imaging and analysis of the transwell assays; immunohistochemical staining, imaging and analysis of the mouse tumor samples. SD2 performed cloning, antibody characterization, analyzed data and assisted with preparing figures. SG performed the *in vivo* analyses including xenograft generation and bioimaging. TTM designed and supervised the *in vivo* analyses. SS prepared recombinant proteins and performed *in vitro* Ni-NTA pull-down experiments. PS, SD1, SD2, and TKK edited the manuscript. TKK conceived and co-ordinated the study and wrote the manuscript. All authors reviewed the results and approved the final version of the manuscript.

## REFERENCES

Amemiya HM, Kundaje A, Boyle AP. 2019. The ENCODE Blacklist: Identification of Problematic Regions of the Genome. Sci Rep 9: 9354.

Arif M, Vedamurthy BM, Choudhari R, Ostwal YB, Mantelingu K, Kodaganur GS, Kundu TK. 2010. Nitric Oxide-Mediated Histone Hyperacetylation in Oral Cancer: Target for a Water-Soluble HAT Inhibitor, CTK7A. Chemistry & Biology 17: 903–913.

Burley SK, Roeder RG. 1996. Biochemistry and structural biology of transcription factor IID (TFIID). Annu Rev Biochem 65: 769–799.

Cho WK, Spille JH, Hecht M, Lee C, Li C, Grube V, Cisse, II. 2018. Mediator and RNA polymerase II clusters associate in transcription-dependent condensates. Science 361: 412–415.

Colombo E, Alcalay M, Pelicci PG. 2011. Nucleophosmin and its complex network: a possible therapeutic target in hematological diseases. Oncogene doi:onc2010646 [pii] 10.1038/onc.2010.646.

Colombo E, Marine JC, Danovi D, Falini B, Pelicci PG. 2002. Nucleophosmin regulates the stability and transcriptional activity of p53. Nat Cell Biol 4: 529–533.

Cong R, Das S, Ugrinova I, Kumar S, Mongelard F, Wong J, Bouvet P. 2012. Interaction of nucleolin with ribosomal RNA genes and its role in RNA polymerase I transcription. Nucleic Acids Res 40: 9441–9454.

Consortium EP. 2012. An integrated encyclopedia of DNA elements in the human genome. Nature 489: 57–74.

Das S, Cong R, Shandilya J, Senapati P, Moindrot B, Monier K, Delage H, Mongelard F, Kumar S, Kundu TK et al. 2013. Characterization of nucleolin K88 acetylation defines a new pool of nucleolin colocalizing with pre-mRNA splicing factors. FEBS Lett 587: 417–424.

Dhar SK, Lynn BC, Daosukho C, St Clair DK. 2004. Identification of nucleophosmin as an NF-kappaB co-activator for the induction of the human SOD2 gene. J Biol Chem 279: 28209–28219.

Dutta S, Akey IV, Dingwall C, Hartman KL, Laue T, Nolte RT, Head JF, Akey CW. 2001. The crystal structure of nucleoplasmin-core: implications for histone binding and nucleosome assembly. Mol Cell 8: 841–853.

Foltz DR, Jansen LE, Black BE, Bailey AO, Yates JR, Cleveland DW. 2006. The human CENP-A centromeric nucleosome-associated complex. Nat Cell Biol 8: 458–469.

Formosa T. 2012. The role of FACT in making and breaking nucleosomes. Biochim Biophys Acta 1819: 247–255.

Frehlick LJ, Eirín-López JM, Ausió J. 2007. New insights into the nucleophosmin/nucleoplasmin family of nuclear chaperones. Bioessays 29: 49–59.

Grisendi S, Mecucci C, Falini B, Pandolfi PP. 2006. Nucleophosmin and cancer. Nat Rev Cancer 6: 493–505.

Grunberg S, Hahn S. 2013. Structural insights into transcription initiation by RNA polymerase II. Trends Biochem Sci 38: 603–611.

Gurard-Levin ZA, Quivy JP, Almouzni G. 2014. Histone chaperones: assisting histone traffic and nucleosome dynamics. Annu Rev Biochem 83: 487–517.

Heinz S, Benner C, Spann N, Bertolino E, Lin YC, Laslo P, Cheng JX, Murre C, Singh H, Glass CK. 2010. Simple combinations of lineage-determining transcription factors prime cis-regulatory elements required for macrophage and B cell identities. Mol Cell 38: 576–589.

Hingorani K, Szebeni A, Olson MO. 2000. Mapping the functional domains of nucleolar protein B23. J Biol Chem 275: 24451–24457.

Hoffman MM, Ernst J, Wilder SP, Kundaje A, Harris RS, Libbrecht M, Giardine B, Ellenbogen PM, Bilmes JA, Birney E et al. 2013. Integrative annotation of chromatin elements from ENCODE data. Nucleic Acids Res 41: 827–841.

Inouye CJ, Seto E. 1994. Relief of YY1-induced transcriptional repression by protein-protein interaction with the nucleolar phosphoprotein B23. J Biol Chem 269: 6506–6510.

Jian Y, Gao Z, Sun J, Shen Q, Feng F, Jing Y, Yang C. 2009. RNA aptamers interfering with nucleophosmin oligomerization induce apoptosis of cancer cells. Oncogene 28: 4201–4211.

Kaplan CD, Laprade L, Winston F. 2003. Transcription elongation factors repress transcription initiation from cryptic sites. Science 301: 1096–1099.

Kaypee S, Sahadevan SA, Sudarshan D, Halder Sinha S, Patil S, Senapati P, Kodaganur GS, Mohiyuddin A, Dasgupta D, Kundu TK. 2018. Oligomers of human histone chaperone NPM1 alter p300/KAT3B folding to induce autoacetylation. Biochim Biophys Acta Gen Subj 1862: 1729–1741.

Kent WJ, Sugnet CW, Furey TS, Roskin KM, Pringle TH, Zahler AM, Haussler D. 2002. The human genome browser at UCSC. Genome Res 12: 996–1006.

Kim D, Langmead B, Salzberg SL. 2015. HISAT: a fast spliced aligner with low memory requirements. Nat Methods 12: 357–360.

Kondo T, Minamino N, Nagamura-Inoue T, Matsumoto M, Taniguchi T, Tanaka N. 1997. Identification and characterization of nucleophosmin/B23/numatrin which binds the anti-oncogenic transcription factor IRF-1 and manifests oncogenic activity. Oncogene 15: 1275–1281.

Langmead B, Salzberg SL. 2012. Fast gapped-read alignment with Bowtie 2. Nat Methods 9: 357–359.

Lee HH, Kim HS, Kang JY, Lee BI, Ha JY, Yoon HJ, Lim SO, Jung G, Suh SW. 2007. Crystal structure of human nucleophosmin-core reveals plasticity of the pentamer-pentamer interface. Proteins 69: 672–678.

Lee TI, Young RA. 2000. Transcription of eukaryotic protein-coding genes. Annu Rev Genet 34: 77–137.

Li B, Carey M, Workman JL. 2007. The role of chromatin during transcription. Cell 128: 707–719.

Li Z, Boone D, Hann SR. 2008. Nucleophosmin interacts directly with c-Myc and controls c-Myc-induced hyperproliferation and transformation. Proc Natl Acad Sci U S A 105: 18794–18799.

Lindström MS. 2011. NPM1/B23: A Multifunctional Chaperone in Ribosome Biogenesis and Chromatin Remodeling. Biochem Res Int 2011: 195209.

Liu JC, Ferreira CG, Yusufzai T. 2015. Human CHD2 is a chromatin assembly ATPase regulated by its chromo-and DNA-binding domains. J Biol Chem 290: 25–34.

Love MI, Huber W, Anders S. 2014. Moderated estimation of fold change and dispersion for RNA-seq data with DESeq2. Genome Biol 15: 550.

Mason PB, Struhl K. 2003. The FACT complex travels with elongating RNA polymerase II and is important for the fidelity of transcriptional initiation in vivo. Mol Cell Biol 23: 8323–8333.

Michaud GA, Snyder M. 2002. Proteomic approaches for the global analysis of proteins. Biotechniques 33: 1308–1316.

Mitrea DM, Cika JA, Stanley CB, Nourse A, Onuchic PL, Banerjee PR, Phillips AH, Park CG, Deniz AA, Kriwacki RW. 2018. Self-interaction of NPM1 modulates multiple mechanisms of liquid-liquid phase separation. Nat Commun 9: 842.

Murano K, Okuwaki M, Hisaoka M, Nagata K. 2008. Transcription regulation of the rRNA gene by a multifunctional nucleolar protein, B23/nucleophosmin, through its histone chaperone activity. Mol Cell Biol 28: 3114–3126.

Namboodiri VM, Akey IV, Schmidt-Zachmann MS, Head JF, Akey CW. 2004. The structure and function of Xenopus NO38-core, a histone chaperone in the nucleolus. Structure 12: 2149–2160.

Ng HH, Robert F, Young RA, Struhl K. 2002. Genome-wide location and regulated recruitment of the RSC nucleosome-remodeling complex. Genes Dev 16: 806–819.

Okuda M. 2002. The role of nucleophosmin in centrosome duplication. Oncogene 21: 6170–6174.

Okuwaki M, Matsumoto K, Tsujimoto M, Nagata K. 2001. Function of nucleophosmin/B23, a nucleolar acidic protein, as a histone chaperone. FEBS Lett 506: 272–276.

Parelho V, Hadjur S, Spivakov M, Leleu M, Sauer S, Gregson HC, Jarmuz A, Canzonetta C, Webster Z, Nesterova T et al. 2008. Cohesins functionally associate with CTCF on mammalian chromosome arms. Cell 132: 422–433.

Pertea M, Pertea GM, Antonescu CM, Chang TC, Mendell JT, Salzberg SL. 2015. StringTie enables improved reconstruction of a transcriptome from RNA-seq reads. Nat Biotechnol 33: 290–295.

Platonova O, Akey IV, Head JF, Akey CW. 2011. Crystal Structure and Function of Human Nucleoplasmin (Npm2): A Histone Chaperone in Oocytes and Embryos. Biochemistry 50: 8078–8089.

Prinos P, Lacoste MC, Wong J, Bonneau AM, Georges E. 2011. Mutation of cysteine 21 inhibits nucleophosmin/B23 oligomerization and chaperone activity. Int J Biochem Mol Biol 2: 24–30.

Qi W, Shakalya K, Stejskal A, Goldman A, Beeck S, Cooke L, Mahadevan D. 2008. NSC348884, a nucleophosmin inhibitor disrupts oligomer formation and induces apoptosis in human cancer cells. Oncogene 27: 4210–4220.

Quinlan AR, Hall IM. 2010. BEDTools: a flexible suite of utilities for comparing genomic features. Bioinformatics 26: 841–842.

Ramirez F, Dundar F, Diehl S, Gruning BA, Manke T. 2014. deepTools: a flexible platform for exploring deep-sequencing data. Nucleic Acids Res 42: W187–191.

Savkur RS, Olson MO. 1998. Preferential cleavage in pre-ribosomal RNA byprotein B23 endoribonuclease. Nucleic Acids Res 26: 4508–4515.

Schwabish MA, Struhl K. 2006. Asf1 mediates histone eviction and deposition during elongation by RNA polymerase II. Mol Cell 22: 415–422.

Schwabish MA, Struhl K. 2007. The Swi/Snf complex is important for histone eviction during transcriptional activation and RNA polymerase II elongation in vivo. Mol Cell Biol 27: 6987–6995.

Senapati P, Dey S, Sudarshan D, Das S, Kumar M, Kaypee S, Mohiyuddin A, Kodaganur GS, Kundu TK. 2018. Oncogene c-fos and mutant R175H p53 regulate expression of Nucleophosmin implicating cancer manifestation. The FEBS journal doi:10.1111/febs.14625.

Senapati P, Sudarshan D, Gadad SS, Shandilya J, Swaminathan V, Kundu TK. 2015. Methods to study histone chaperone function in nucleosome assembly and chromatin transcription. Methods in molecular biology (Clifton, NJ) 1288: 375–394.

Shandilya J, Senapati P, Hans F, Menoni H, Bouvet P, Dimitrov S, Angelov D, Kundu TK. 2014. Centromeric histone variant CENP-A represses acetylation-dependent chromatin transcription that is relieved by histone chaperone NPM1. Journal of Biochemistry 156: 221–227.

Shandilya J, Swaminathan V, Gadad SS, Choudhari R, Kodaganur GS, Kundu TK. 2009. Acetylated NPM1 Localizes in the Nucleoplasm and Regulates Transcriptional Activation of Genes Implicated in Oral Cancer Manifestation. Molecular and Cellular Biology 29: 5115–5127.

Soutourina J, Bordas-Le Floch V, Gendrel G, Flores A, Ducrot C, Dumay-Odelot H, Soularue P, Navarro F, Cairns BR, Lefebvre O et al. 2006. Rsc4 connects the chromatin remodeler RSC to RNA polymerases. Mol Cell Biol 26: 4920–4933.

Stark C, Breitkreutz BJ, Reguly T, Boucher L, Breitkreutz A, Tyers M. 2006. BioGRID: a general repository for interaction datasets. Nucleic Acids Res 34: D535–539.

Subramanian A, Tamayo P, Mootha VK, Mukherjee S, Ebert BL, Gillette MA, Paulovich A, Pomeroy SL, Golub TR, Lander ES et al. 2005. Gene set enrichment analysis: a knowledge-based approach for interpreting genome-wide expression profiles. Proc Natl Acad Sci U S A 102: 15545–15550.

Swaminathan V, Kishore A, Febitha K, Kundu T. 2005. Human histone chaperone nucleophosmin enhances acetylation-dependent chromatin transcription. Molecular and Cellular Biology 25: 7534–7545.

Szebeni A, Olson MO. 1999. Nucleolar protein B23 has molecular chaperone activities. Protein Sci 8: 905–912.

Wendt KS, Yoshida K, Itoh T, Bando M, Koch B, Schirghuber E, Tsutsumi S, Nagae G, Ishihara K, Mishiro T et al. 2008. Cohesin mediates transcriptional insulation by CCCTC-binding factor. Nature 451: 796–801.

Yusufzai TM, Tagami H, Nakatani Y, Felsenfeld G. 2004. CTCF tethers an insulator to subnuclear sites, suggesting shared insulator mechanisms across species. Mol Cell 13: 291–298.

Zeller KI, Haggerty T, Barrett JF, Guo Q, Wonsey DR, Dang CV. 2001. Characterization of nucleophosmin (B23) as a Myc target by scanning chromatin immunoprecipitation (SChIP). doi:10.1074/jbc.M108506200.

Zhang Y, Liu T, Meyer CA, Eeckhoute J, Johnson DS, Bernstein BE, Nusbaum C, Myers RM, Brown M, Li W et al. 2008. Model-based analysis of ChIP-Seq (MACS). Genome Biol 9: R137.

